# AGO1 in association with *NEAT1* lncRNA contributes to nuclear and 3D chromatin architecture in human cells

**DOI:** 10.1101/525527

**Authors:** Muhammad Shuaib, Krishna Mohan Parsi, Hideya Kawaji, Manjula Thimma, Sabir Abdu Adroub, Alexandre Fort, Yanal Ghosheh, Tomohiro Yamazaki, Taro Mannen, Loqmane Seridi, Bodor Fallatah, Waad Albawardi, Timothy Ravasi, Piero Carninci, Tetsuro Hirose, Valerio Orlando

## Abstract

Aside from their roles in the cytoplasm, RNA-interference components have been reported to localize also in the nucleus of human cells. In particular, AGO1 associates with active chromatin and appears to influence global gene expression. However, the mechanistic aspects remain elusive. Here, we identify AGO1 as a paraspeckle component that in combination with the *NEAT1* lncRNA maintains 3D genome architecture. We demonstrate that AGO1 interacts with *NEAT1* lncRNA and its depletion affects *NEAT1* expression and the formation of paraspeckles. By Hi-C analysis in AGO1 knockdown cells, we observed global changes in chromatin organization, including TADs configuration, and A/B compartment mixing. Consistently, distinct groups of genes located within the differential interacting loci showed altered expression upon AGO1 depletion. *NEAT1* knockout cells displayed similar changes in TADs and higher-order A/B compartmentalization. We propose that AGO1 in association with *NEAT1* lncRNA can act as a scaffold that bridges chromatin and nuclear bodies to regulate genome organization and gene expression in human cells.

## Introduction

Argonaute (AGO) proteins are key components of the RNA interference (RNAi) pathway and are mainly recognized for their role in post-transcriptional gene silencing via RNA guided mechanisms in the cytoplasm (Joshua-Tor and Hannon 2011; Meister 2013). In fission yeast Ago proteins and small RNAs are essential for heterochromatin formation (Volpe et al. 2002) and, similarly, in plants in RNA-dependent DNA methylation (Zilberman et al. 2003). However, a direct function of RNAi components in animal cell chromatin-mediated transcriptional regulation remains unclear. We previously reported that in *Drosophila*, RNAi components (dAgo2 and Dicer) are preferentially associated with active chromatin and affect transcription by regulating RNA Pol II pausing and stress response (Cernilogar et al. 2011). Another report indicated that in *Drosophila* dAgo2 associates with insulator proteins CTCF/CP190 at active promoters and controls CTCF/CP190-dependent looping interactions between insulators, promoters, and enhancers (Moshkovich et al. 2011). In human cells, AGO1 binds RNA pol II (Huang et al. 2013) to associate with active promoters (Huang et al. 2013), enhancers (Allo et al. 2014), HP1α, and CTCF (Agirre et al. 2015) and affects splicing (Ameyar-Zazoua et al. 2012; Allo et al. 2014; Agirre et al. 2015). The association of AGO1 with active enhancers was postulated to be mediated by long nuclear RNAs (Allo et al. 2014). However, the involvement of RNA(s) in AGO1 nuclear function and its direct or indirect mechanistic effects on chromatin and transcription remains unclear.

A body of evidence indicates a functional role of some long non-coding RNAs (lncRNAs) in controlling the activity of chromatin remodelers and nuclear compartmentalization (Rinn and Chang 2012). Among lncRNAs, Nuclear Enriched Assembly Transcript 1 (*NEAT1*) is an essential structural component of nuclear bodies called paraspeckles. The *NEAT1* locus produces two isoforms of *NEAT1* transcripts (3.7-kb *NEAT1_1* and 23-kb *NEAT1_2* in human), and its transcription is the first step in paraspeckle construction (Clemson et al. 2009; Mao et al. 2011). The concomitant association with >50 “paraspeckle associated” proteins (PSPs) produces a *de novo* paraspeckle (Sasaki et al. 2009; Naganuma et al. 2012; Chujo et al. 2017). Though the exact function of paraspeckles remains unclear, their dynamic assembly is believed to regulate gene expression (Prasanth et al. 2005; Hirose et al. 2014; Imamura et al. 2014). The genome-wide association of *NEAT1* lncRNA with active sites (West et al. 2014) and phase-separation mediated *NEAT1* dependent paraspeckles assembly (Yamazaki et al. 2018) suggest that *NEAT1* paraspeckle and its associated protein factors may be required for regulating nuclear architecture and gene expression. However, a potential contribution of the *NEAT1* lncRNA and paraspeckles in shaping three-dimensional chromatin organization has not yet been tested.

Recently, genome-wide chromosome conformation capture analysis (Hi-C) has revealed that chromosomes are organized into active (open, A-type) and inactive (closed, B-type) compartments (Lieberman-Aiden et al. 2009), which are further composed of clusters of interactions called Topologically Associated Domains (TADs), separated by boundary regions (Dixon et al. 2012; Nora et al. 2012). The 3D chromatin structure is believed to regulate gene expression by establishing loops between enhancers and promoters, and/or by bridging regulatory elements and genes into spatial chromatin hubs, compartments, and domains (Bouwman and de Laat 2015). Several chromatin regulators, including Polycomb proteins, Cohesin, CTCF, and Mediator complex have been reported to influence 3D genome organization (Kagey et al. 2010; Delest et al. 2012; Seitan et al. 2013; Sofueva et al. 2013; Zuin et al. 2014). However, the collective role of RNA binding proteins like RNAi components and lncRNA (Yang et al. 2013) has been poorly investigated.

Here, we have applied a combination of genome-wide approaches to investigate the role of chromatin-bound AGO1 and its associated RNA in 3D genome conformation and gene expression. Collectively, our data unveil an unprecedented link between nuclear AGO1 and *NEAT1* lncRNA, which contribute to the stability of sub-nuclear compartment and fine-tuning of higher-order chromatin architecture and gene expression in human cells.

## Results

### AGO1 depletion results in deregulation of coding and non-coding transcripts at the genome-wide level

Despite the previously reported association of AGO1 with active promoters and enhancers, its direct or indirect effects on transcription remain unclear. To evaluate the impact of AGO1 depletion on global transcriptional changes of all expressed TSS, we decided to use CAGE-seq analysis (Takahashi et al. 2012). We applied CAGE-seq analysis on total, and chromatin extracted RNA from control (siCtrl) and AGO1 depleted (siAGO1) cells (Supplemental Fig. S1e-i, and see supplemental results). We identified more than 1,000 genes in both total and chromatin-bound RNA CAGE-seq data that were differentially expressed (DE) (Fig. 1A-B and Supplemental Fig. S2a-b). We observed deregulation of not only coding transcripts but also a significant change in noncoding transcript levels (Fig. 1C). These observations indicate that loss of AGO1 has both a positive and negative impact on gene expression (Supplemental Fig. S2c). We further extended our analysis to find a possible correlation between perturbed transcripts and deregulated miRNAs. To this, we compared small RNA-seq data (Supplemental Fig. S2d) in control, and AGO1 depleted cells. The majority (95%) of perturbed mRNAs were not direct targets of differentially expressed miRNAs (Supplemental Fig. S2e). Moreover, we did not observe any significant correlation between the effect of down- and up-regulated miRNAs and the expression of predicted targets (Supplemental Fig. S2f-i). To discriminate between directly and indirectly regulated genes, we integrated CAGE-seq with AGO1 binding sites on chromatin (ChIP-seq). First, we examined AGO1 genomic distribution by ChIP-seq in HepG2 cells (Supplemental Fig. S3a-b, and see supplemental results). As previously reported (Allo et al. 2014) we found AGO1 enrichment across many active sites such as enhancers, promoters and also other genic regions (Supplemental Fig. S3c-d). Among all the differentially expressed genes, we observed AGO1 binding at about 29% (274) genes, but only 16% (43) of these genes overlapped with AGO1 peaks at their promoters. In contrast, for the majority of perturbed genes (71%) we did not detect AGO1 at their promoters or within the coding region (Fig. 1D). However, we observed a clear enrichment of AGO1 binding at the predicted enhancers located within <50 kb of all the differentially expressed genes (Fig. 1E). We further categorized AGO1 peaks into three classes, i.e., promoter, proximal and distal regions and observed a higher enrichment at distal sites (Fig. 1F and Supplemental Fig. S3e). As expected, these AGO1 enriched regions showed high overlap with ENCODE HepG2 histone ChIP-seq datasets such as H3K4me1, H3K4me3, H3K27ac, and H3K9ac, representing active enhancers (Supplemental Fig. S3f). Next, by overlapping all transcripts from CAGE-seq with AGO1 peaks, we observed a strong correlation with RNAs produced at those sites (Fig. 1G). We found detectable levels of transcriptional activity at AGO1 bound proximal and distal regions, which represent putative enhancer RNAs (eRNA) (Fig. 1G), We also performed AGO2 ChIP-seq with a specific antibody, but we failed to obtain enough reproducible peaks (data not shown), which is consistent with a previous report (Huang et al. 2013). Furthermore, cellular fractionation followed by western-blot analysis revealed higher AGO1 enrichment compared to AGO2 and AGO3 in the chromatin-bound fraction (Supplemental Fig. S1b). Thus, the conserved association of AGO1 with active chromatin sites (Cernilogar et al. 2011; Huang et al. 2013) and the lack of its endonuclease activity, but not the RNA binding domain (Wu et al. 2008) suggested an RNA processing independent role of AGO1 in chromatin. Therefore, we focused our subsequent analysis only on AGO1 and its potential interacting RNA partners.

**Figure 1.**
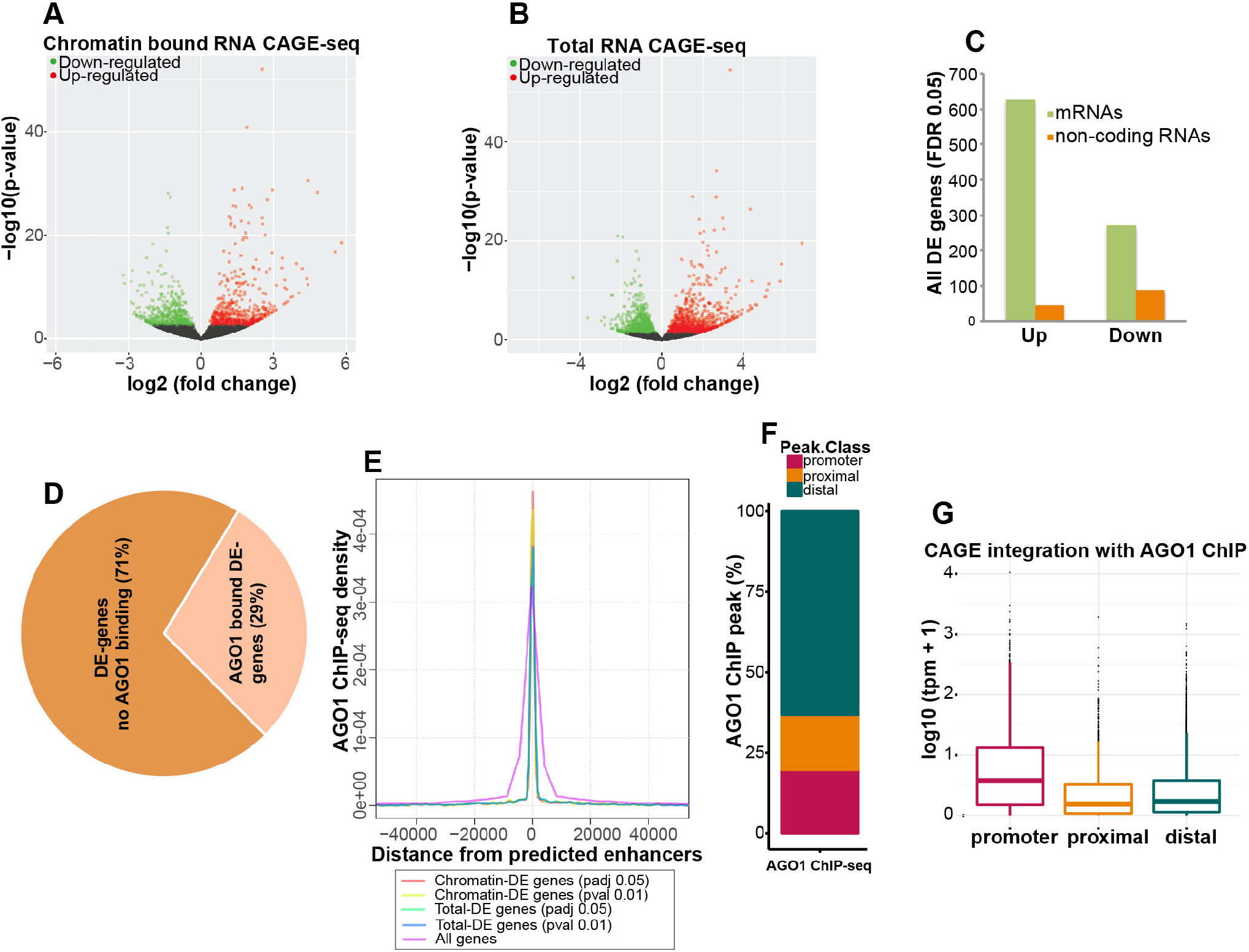
AGO1 knockdown perturbs expression of coding and non-coding transcripts. (*A-B*) Volcano plots showing differentially expressed genes in AGO1 depleted cells, identified by CAGE-seq analysis from (*A*) chromatin associated and (*B*) total RNA. Significantly down-regulated (green) and up-regulated (red) genes are shown. (*C*) Total and chromatin associated differentially expressed coding (mRNA) and noncoding genes (FDR 0.05). (*D*) Integration of ChIP-seq and AGO1-knockdown CAGE-seq data classify AGO1-responsive genes into direct and indirect targets. 29% of all differentially expressed (DE) genes (from total and chromatin CAGE-seq combined) contain AGO1 binding at their promoter or intragenically, represent direct targets. (*E*) Density plot of AGO1 ChIP-seq peaks closer than 50-kb to genes (either differentially expressed or considering all genes) centered at enhancer regions predicted by ChromHMM. (*F*) Bar plot shows percent enrichment of AGO1 ChIP-seq peaks at distal, promoter, and proximal regions. AGO1 peaks are highly enriched in distal regions (i.e, > 5-kb from nearest annotated TSS). A significant amount of AGO1 peaks also associate with promoter region (i.e, < 1-kb from nearest annotated TSS). (*G*) Density of CAGE-seq signals represents transcriptional activity at AGO1 binding regions (promoter, proximal and distal). The low level of transcriptional activity at distal regions indicates enhancer and lncRNAs.

### Analysis of chromatin-bound AGO1 associated RNAs identifies *NEAT1* lncRNA as a highly enriched component

To clarify the involvement of RNA(s) in AGO1 nuclear function, we explored chromatin-bound AGO1-associated RNA moieties in the nucleus by cross-linked RIP-seq. For maximum output of RIP-seq data, the extracted RNA molecules were separated into two fractions (~100 nt fraction and ~400 nt fraction) and processed for deep sequencing using the Illumina platform. The two fractions of AGO1 RIP-seq data displayed similar profiles (Fig. 2A). The genome-wide mapping of the AGO1-associated RNA fractions (~100 and ~400 bp) showed enrichment of coding, non-coding and un-annotated RNA peaks (Fig. 2B-C). Additionally, we intersected AGO1-associated RNA peaks with AGO1-bound (ChIP-seq) and -unbound chromosomal regions across the genome. They showed a significant enrichment on AGO1-bound active sites (promoters and enhancers) when compared to AGO1-absent sites in the genome (Fig. 2D). Next, we identified a total of 3500 and 1854 transcripts containing approximately 50% non-coding RNAs in both 100 bp and 400 bp RIP-seq fractions respectively.

**Figure 2.**
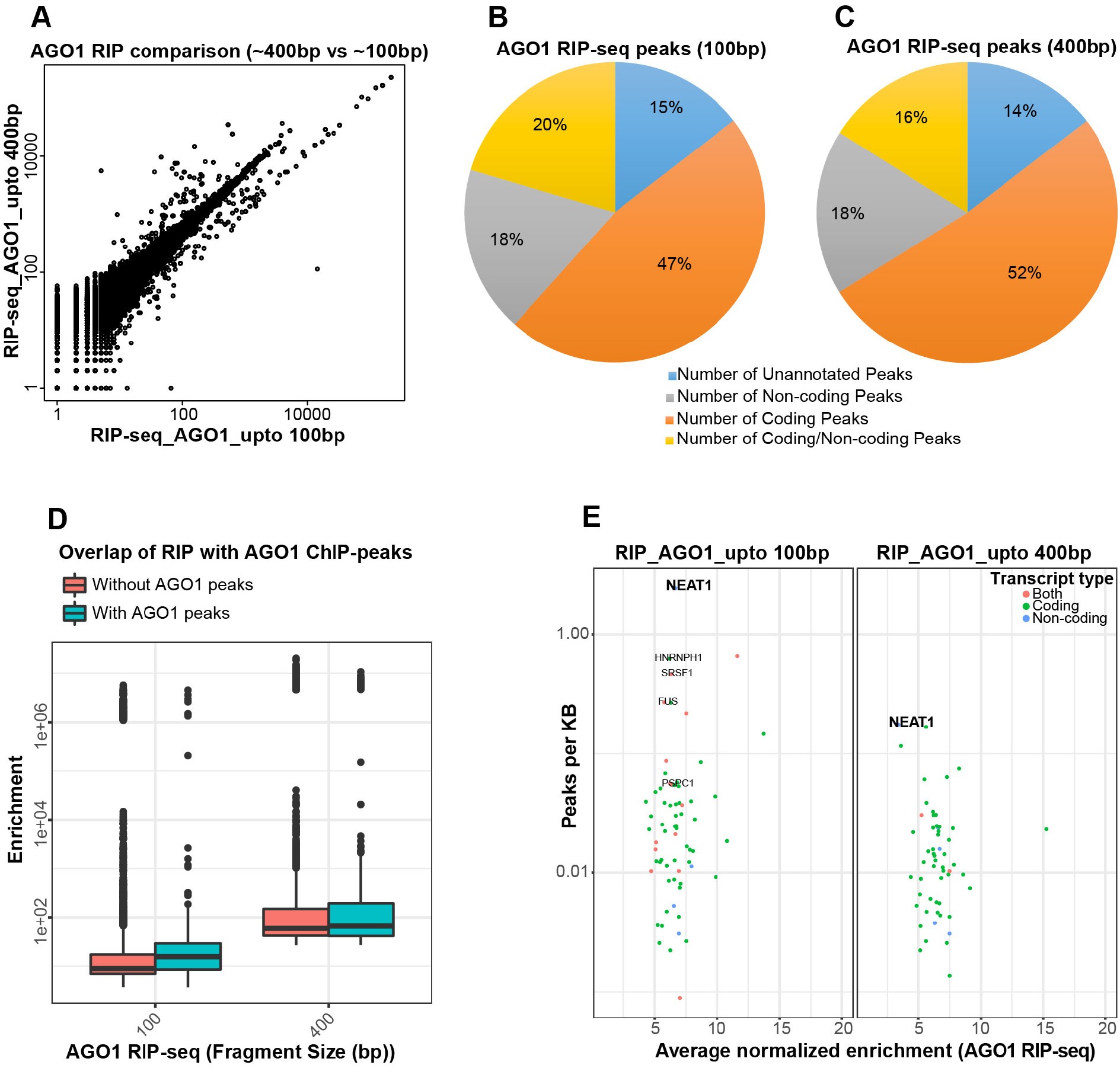
Identification of chromatin-bound AGO1-associated RNAs. (*A*) We performed RIP-seq of two RNA fractions (~100 bp and ~400 bp) bound to chromatin-extracted AGO1. Mapping sequencing reads from these two fractions display similar distributions (*B-C*) Pie-charts show AGO1 RIP-seq peaks classification identified after mapping and peak-calling using Piranha (V1.2.1) in the two fractions: (*B*) ~100 bp (*C*) ~400 bp. (*D*) The Box-plot shows enrichment of RIP-seq peaks (both 400bp and 100bp fractions) on AGO1 binding sites (ChIP-seq peaks) (Mann-Whitney test, p-value 4.4e-12) (*E*) Representation of highly enriched AGO1 associated chromatin bound transcripts (coding, non-coding and both).

Among AGO1-associated lncRNAs, we identified *NEAT1* lncRNA as the most significantly enriched major partner of chromatin-bound AGO1 in both RNA fractions (Fig. 2E). *NEAT1* lncRNA is an essential structural component of paraspeckle nuclear bodies. Along with *NEAT1*, we also found enriched transcripts of other paraspeckle components such as *PSPC1, HNRNPH1* and *FUS* (Fig. 2E). These results suggested that the nuclear AGO1 associates with paraspeckles and its major component *NEAT1* lncRNA.

### AGO1 interacts with *NEAT1* and other essential paraspeckle proteins and is required for paraspeckle integrity

Given the enrichment of *NEAT1* in AGO1-bound chromatin fraction, we sought to verify whether AGO1 is an integral component of paraspeckle nuclear bodies. Along with *NEAT1* short isoform (3.7-kb), we found a higher number of reads coverage on *NEAT1_2* (23-kb) (Fig. 3A), which is essential for *de novo* assembly of paraspeckles. The direct interaction between AGO1 and both *NEAT1* isoforms (*NEAT1_1* and *NEAT1_2*) was confirmed by UV cross-linking and immunoprecipitation followed by qPCR (Fig. 3B and Supplemental Table S1). Immunofluorescence combined with RNA FISH revealed AGO1 co-localization with *NEAT1* lncRNA in paraspeckles (Fig. 3C), but not AGO2 (Supplemental Fig. S1d). The specificity of AGO1 detection was confirmed by observing signal disappearance upon AGO1 depletion (Supplemental Fig. S1i). Also, we used soluble nuclear, and chromatin extracts of stable cell lines expressing AGO1 and AGO2 fused with C-terminal Flag- and HA-epitope tags (named e-AGO1 and e-AGO2) for double-immunoaffinity purification (Supplemental Fig. S4a). The essential paraspeckle proteins were identified explicitly in the e-AGO1 but not e-AGO2 nuclear and chromatin complexes by mass spectrometry and western blotting (Fig. 3D-E and Supplemental Fig. S4b). We next conducted *NEAT1* ChIRP-western blot experiment to confirm the interaction of AGO1 with *NEAT1* (Supplemental Fig. S5a-b). *NEAT1* ChIRP followed by immunoblotting further proved the presence of AGO1 along with paraspeckle proteins (SFPQ and NONO) in the *NEAT1* associated proteins (Supplemental Fig. S5c-d). These results indicate that AGO1 is stably associated with paraspeckle.

**Figure 3.**
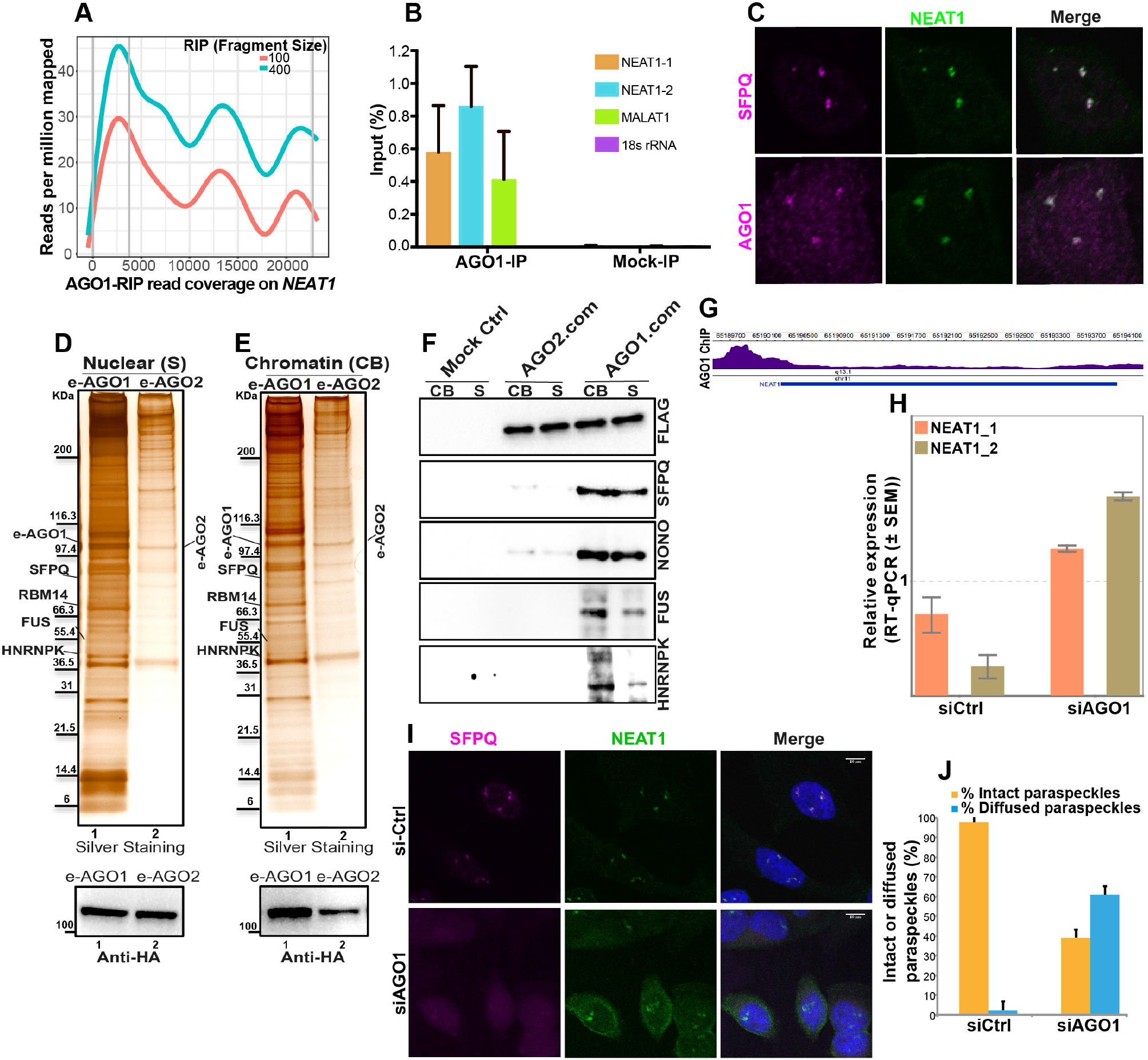
AGO1 co-localizes and interacts with paraspeckle components, and its knockdown impact paraspeckles formation. (*A*) Read coverage enrichment on both short and long isoforms of *NEAT1* lncRNA in AGO1 RIP-seq fraction (100 bp and 400 bp). (*B*) Validation of direct interaction between AGO1 and *NEAT1* lncRNA by UV crosslinking IP. After UV cross-linking, the RNAs immunoprecipitated with AGO1 IP were quantified by qRT-PCR, and the RNA enrichment (as % input) was determined (± SD from 3 experiments). IgG IP was used as mock control. (*C*) Localization of AGO1 and SFPQ, a known interactor, relative to paraspeckles in human HepG2 cells. AGO1 and SFPQ were detected by immunofluorescence and *NEAT1* RNA-FISH was used to visualize paraspeckles. (*D-F*) Immunopurification of FALG-HA epitope tagged e-AGO1 and e-AGO2 complexes from soluble nuclear and chromatin fractions. (*D-E*) Silver staining (top) of e-AGO1 (lane1) and e-AGO2 (lane2) associated proteins from (*D*) nuclear soluble and (*E*) chromatin fractions. The protein complexes containing e-AGO1 and e-AGO2 were purified by double immunoaffinity from soluble nuclear and chromatin extracts. Protein bands were identified by mass spectrometry analysis and the positions of molecular size markers are indicated. Paraspeckle components were identified in e-AGO1 nuclear and chromatin complexes. (*D-E*) Western blot (lower) using HA antibody shows level of e-AGO1 (lane1) and e-AGO2 (lane2) in (*D*) nuclear soluble and (*E*) chromatin complexes. (*F*) The e-AGO1, e-AGO2 and mock control (Mock ctrl) nuclear soluble (S) and chromatin (CB) complexes (AGO1-com and AGO2-com) were analyzed by immunoblotting with the indicated antibodies. Essential paraspeckle proteins SFPQ, NONO, FUS, and HNRNPK are specific to e-AGO1 nuclear and chromatin complex. Double immunoaffinity was performed on untagged HEK-293 nuclear and chromatin extract as a mock control (Mock ctrl). (*G*) Enrichment of AGO1 ChIP-seq signal (purple) on the promoter locus (chr11) of *NEAT1* lncRNA. (H) RT-qPCR analysis of *NEAT1_1* and *NEAT1_2* expression in siCtrl and siAGO1 HepG2 cells. Values were normalized relative to the geometric mean of 18S rRNA, GAPDH, and Actin-B mRNA expression levels. Error bars indicate mean ± s.e.m. (*I*) Examining the effects of AGO1 depletion on paraspeckle formation. Paraspeckle integrity was examined by SFPQ localization to *NEAT1* paraspeckles using a combination of RNA-FISH and immunofluorescence. (*J*) Quantification of the number of cells having an intact or diffuse paraspeckle signal as shown in (*I*). Bar graph represents mean (± s.e.m) of three technical replicates. We scored paraspeckle morphology in 670 cells (for the combined replicates) of siCtrl and siAGO1 HepG2.

Our AGO1 ChIP-seq peaks showed enrichment at the promoter of the *NEAT1* gene (Fig. 3G), indicating the *NEAT1* locus to be a direct target of AGO1. Indeed, upon AGO1 depletion by CAGE-seq, we observed up-regulation of the *NEAT1* transcript. Importantly, by RT-qPCR we confirmed the induction of both *NEAT1_1* and *NEAT1_2*) in AGO1 knockdown cells (Fig. 3H). The expression of *NEAT1* is an essential early step in paraspeckle formation, thus we asked whether AGO1 is involved in paraspeckle assembly. We examined the paraspeckle integrity after AGO1 knockdown by analyzing co-localization of *NEAT1* with SFPQ an essential paraspeckle component. Despite increased *NEAT1* RNA-FISH signal, we observed a diffuse SFPQ signal in >60% of siAGO1 cells, affecting paraspeckle integrity (Fig. 3I-J). Furthermore, by ChIRP-western blot we checked the effect of AGO1 depletion on the interaction of other essential paraspeckle proteins with *NEAT1*. In AGO1 depleted cells, immunoblot analyses of *NEAT1* associated proteins showed a reduction in the level of crucial paraspeckle components (NONO, HNRNPK, and SFPQ) (Supplemental Fig. S5e). In summary, AGO1 interacts with *NEAT1* lncRNA and other paraspeckle components, and its depletion impacts *NEAT1* expression and formation of intact paraspeckles.

### AGO1 depletion alters chromatin interactions specifically in AGO1 bound regions

Since AGO1 binds specific gene loci and it is necessary for intact paraspeckle formation, we sought to investigate the role of AGO1 in higher order chromatin organization. To this end, we performed an unbiased genome-wide chromatin conformation analysis using Hi-C in siCtrl and siAGO1 HepG2 cells (Supplemental Fig. S6). At the global level, the chromosomal interaction frequency heatmaps for each chromosome displayed similar patterns in both siAGO1 and siCtrl cells (Fig. 4A-B). To gain deeper insight into differential interaction frequencies between siCtrl and siAGO1 cells, we compared normalized HiC data (siAGO1 versus siCtrl) with the R package diffHiC (Lun and Smyth 2015). In this case, significant differences in genome-wide chromatin interactions between siAGO1 and siCtrl cells were revealed (Fig. 4C). Upon AGO1 depletion, we observed extensive changes in both long-range and short-range interaction frequency across the genome (Fig. 4C and Supplemental Fig. S7a). Significant differences were detected in siAGO1 cells in both intra- and inter-chromosomal interactions (Fig. 4D and Supplemental Fig. S7b-c). Next, we asked whether changes in chromosomal interactions were correlated with AGO1 binding sites. We found that 93% of AGO1 binding sites (16,566 AGO1 peaks) overlapped with differential interacting regions at a genome-wide level (Fig. 4E). We then calculated the difference in the number of differential interactions between AGO1 enriched and without AGO1 binding 20-kb bins. We observed a significantly (t-test < 0.05) higher number of differential interactions between bin-pairs containing AGO1 binding sites than without AGO1 binding (Fig. 4F). In particular, upon AGO1 depletion we observed a significant decrease in chromosomal interaction frequency among AGO1 binding sites, with a simultaneous gain in chromatin interactions at other genomic loci (Fig. 4G). Overall, these data suggest that AGO1 binding at genomic regulatory sites influences higher order chromatin interactions.

**Figure 4.**
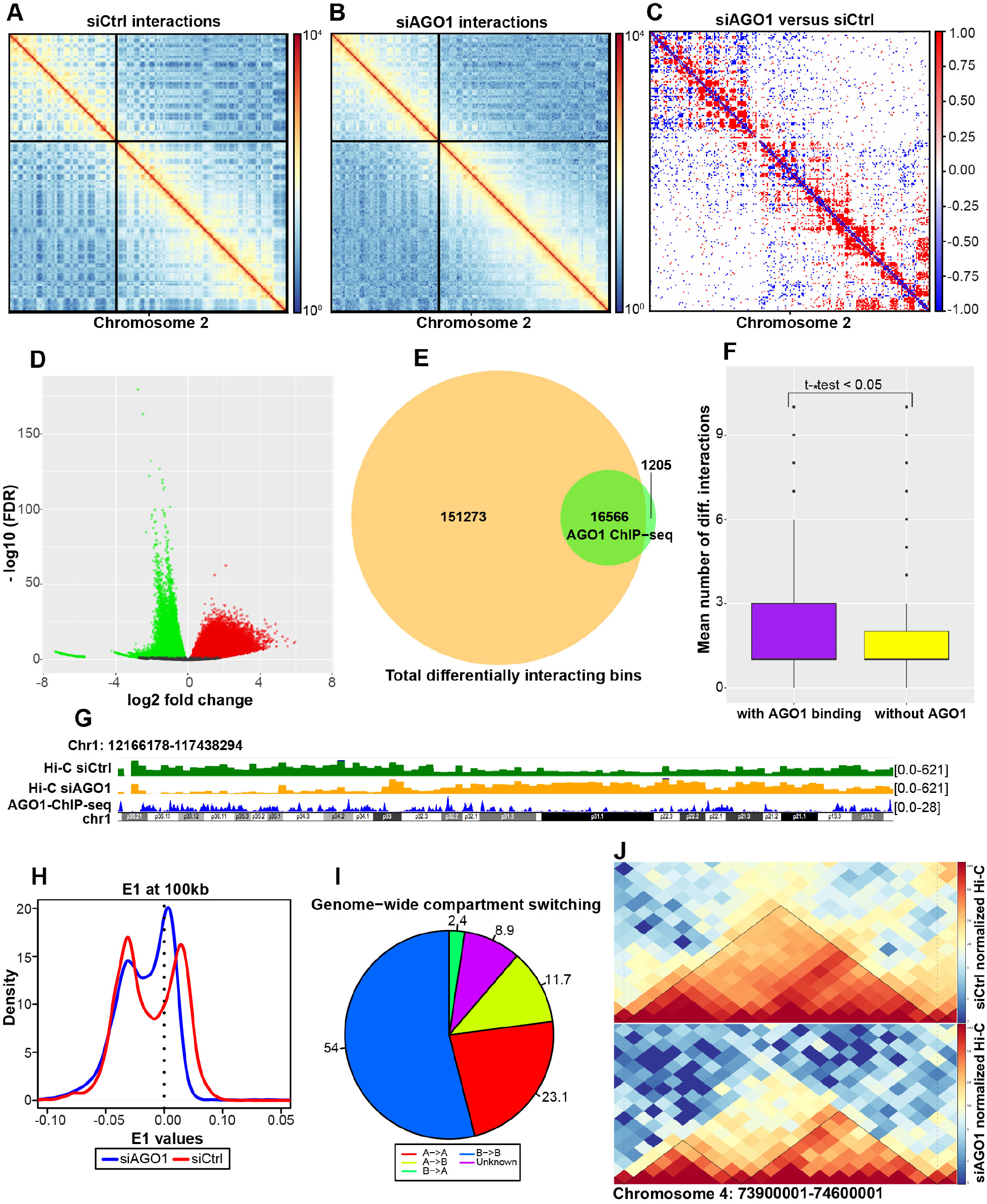
Changes in chromatin interactions, compartments, and TADs upon AGO1 knockdown. Genome-wide chromatin conformation Hi-C analysis was performed using two replicates of siCtrl and siAGO1 cells. Representative normalized Hi-C interaction heatmaps of Chromosome 2 at 1Mb resolution are shown in (*A*) siCtrl and (*B*) siAGO1 cells. (*C*) Differential interaction heatmap for chromosome 2 (1Mb), showing up-regulated (red) and down-regulated (blue) interacting bins. (*D*) Volcano plot representing differential interactions (DI) at 20 kb resolution. The up-regulated interactions are shown in red and the down-regulated interactions are shown in green. (*E*) Venn diagrams showing the number of AGO1 ChIP-seq peaks overlapping with all differential interacting bins, at 20-kb resolution. (*F*) Plot showing mean number of differential interactions calculated in 20-kb bins, based on the presence and absence of overlapping AGO1 peaks at genome-wide level (t-test, p-value < 0.05 and Montecarlo permutation test (999): p-value ≤ 0.002). (*G*) Genome browser snapshot showing Hi-C interaction frequency in siCtrl (green track), siAGO1 (orange track), and AGO1 ChIP-seq peaks (blue track) at chromosome 1 locus: 12166178-117438294. (*H*) Density plot showing the distribution of eigenvalues of siCtrl and siAGO1 cells at 100kb resolution. siCtrl cells followed a bimodal distribution but not the siAGO1 cells. (*I*) Pie chart showing chromatin compartment changes between siCtrl and siAGO1 genomes. “A” and “B” represent open (active) and closed (inactive) compartments, respectively. “A->A” means no change in the compartment type between siCtrl and siAGO1 cells. “B−>B” means no change in the compartment type between siCtrl and siAGO1 cells. “A−>B” represents compartments that are A-type in siCtrl but changed to B-type in siAGO1. “B−>A” represents compartments that are B-type in siCtrl but changed to A-type in siAGO1. Unknown represents when the compartment status is not known in either siCtrl or siAGO1. (*J*) Visualization of normalized Hi-C interactions from siCtrl and siAGO1 cells as twodimensional heat-maps showing TADs in both conditions. TADs found in siCtrl cells are often subdivided into two or more TADs in siAGO1 cells. An example from Chromosome 4 (locus: 73900001-74600001) is shown.

### AGO1 depletion perturbs chromatin compartments and TADs organization

Globally, the genome is partitioned into active (open, A-type) and inactive (closed, B-type) compartments (Lieberman-Aiden et al. 2009; Rao et al. 2014). Thus, we first investigated the effects of AGO1 depletion on the A/B chromatin compartmentalization. We defined chromatin compartments (A and B) using the first principal component (PC1) by eigenvector analysis (Zhang et al. 2012). To further demarcate A/B chromatin compartments, we also intersected expressed genes (identified by CAGE-seq) with eigenvector values. We then compared the genome-wide compartments between siCtrl and siAGO1 cells at 100 kb resolution. We observed changes in the distribution of open and closed compartments in siAGO1 compared to siCtrl cells (Fig. 4H). Globally, the siAGO1 displayed a highly mixed A and B compartmentalization (Fig. 4H and Supplemental Fig. S8a). Notably, 24% of the siAGO1 genome showed compartments switching, which predominantly occurred in the open A-type compartment (Fig. 4I).

To examine the effect on TAD organization, we first identified 3,143 and 3,593 TADs in siCtrl and siAGO1 cells respectively using TADbit tool (Serra 2016). We consistently found that in siAGO1 TADs size was smaller compared to siCtrl cells (t-test, FDR < 0.05) (Supplemental Fig. S8b). Then, by visualizing the topological domains in AGO1-depleted cells, we frequently observed that larger domains found in siCtrl cells were divided into smaller sub-domains (Fig. 4J). Additionally, we found that 25% of AGO1 binding sites (detected by ChIP-seq) overlapped with TAD boundaries identified at 100-kb resolution (Supplemental Fig. S8c). Finally, although there are differences in the numbers and size of TADs in siAGO1 cells, TAD boundaries between siCtrl and siAGO1 cells were largely maintained (Supplemental Fig. S8d). These results suggest that AGO1 contributes to the maintenance of TADs and chromatin compartmentalization.

### AGO1 dependent topological changes correlate with differential gene expression

To determine whether AGO1 dependent changes in chromatin topological structure correlated with differential gene expression, we applied an integrative approach by combining transcriptome (CAGE-seq) with Hi-C and AGO1 ChIP-seq analyses. We first examined the effect of changes in compartmentalization on gene expression. We found that in siAGO1 cells 20% of differentially expressed genes were associated with disorganized chromatin compartments (Fig. 5A-C). Notably, all deregulated genes were present in the active (A) compartments in wild-type cells. Furthermore, the majority of deregulated genes that reside within topological domains, or near domain borders, overlapped with extensive changes in chromosomal interactions following AGO1 knockdown (Fig. 5D). Next, we classified all deregulated genes (FDR ≤ 0.05 and log2FC >1.2) located within differential interacting bins (DI bin) into two categories based on correlation with either i) compartment switching or ii) changes in TADs configuration (Fig. 5E). We further validated by RT-qPCR, the changes in expression of selected perturbed genes (AGO1 KD CAGE-seq) located within differentially interacting bins (AGO1 KD Hi-C) (Supplemental Fig. S9a-d). Altogether, around 80% of deregulated genes are correlated with differential interactions, and a subset of these genes is affected by compartment transitions upon AGO1 depletion.

**Figure 5.**
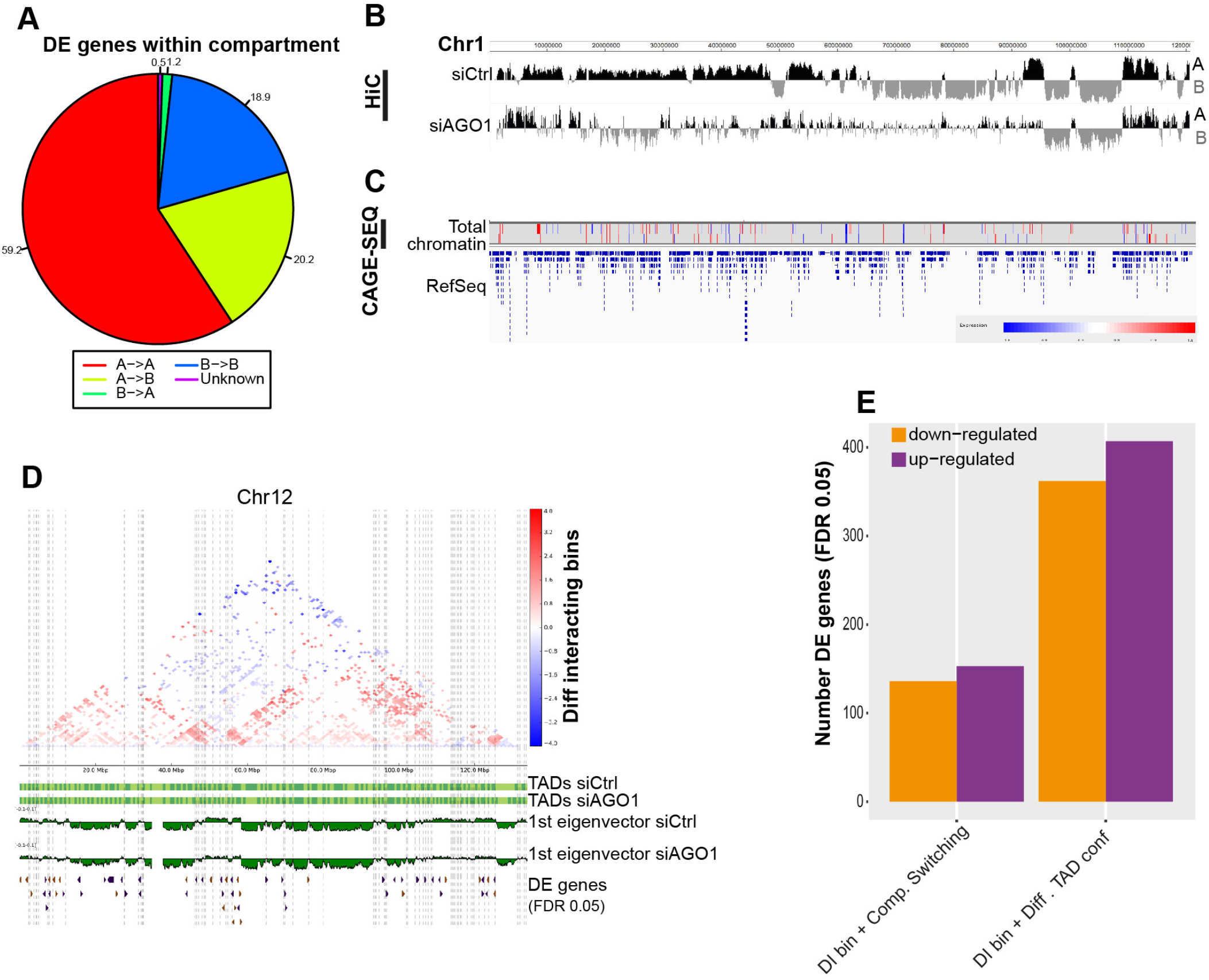
Integration of changes in chromatin topology with differential gene expression after AGO1-depletion. (*A*) Pie chart represents distribution of all differentially expressed (DE) genes (identified by CAGE-seq upon AGO1 depletion) in A/B compartments between siCtrl and siAGO1 cells. “A->A” and “B->B” represent DE genes that remain in the same “A” or “B” compartments respectively. “A->B” represents DE genes that switched from “A” to “B” compartment and “B->A” represents DE genes that switched from “B” to “A” compartment in siAGO1 cells. (*B*) Visualization of chromatin compartments A/B showing changes between siCtrl and siAGO1 cells (chromosome 1). (*C*) Genome browser snapshot showing differentially expressed genes in the same region of chromosome 1 as in (b), up-regulated in red and down-regulated in blue (threshold FDR 0.05). (*D*) A representative plot of Chromosome 12) showing i. differential interactions up-interacting bins (red) and down-interacting bins (blue) ii. TADs configuration in siCtrl and siAGO1 cells (100-kb resolution) iii. Compartments in siCtrl and siAGO1 (100-kb resolution). The vertical dotted lines reflect compartment switching between siCtrl and siAGO1 cells. iv. Differentially expressed genes with up-regulated in purple and down-regulated in brown (threshold FDR 0.05) (*E*) Bar plot showing classification of differentially expressed genes into two categories based on integration with AGO1 ChIP-seq and Hi-C data (TADs and compartment switching were called at 100-kb, and differentially interacting bins at 20-kb resolution). This plot contains the number of DE genes (up-regulated and down-regulated, with FDR threshold 0.05) overlapping with AGO1-bound differentially interacting bins that either associate with different TADs configuration, and or compartment switching between siCtrl and siAGO1 cells.

### *NEAT1* depletion impacts AGO1 paraspeckle localization and chromatin architecture

To understand the functional link between *NEAT1* and AGO1, we asked whether *NEAT1* is necessary and sufficient for AGO1 nuclear paraspeckle localization. First, we examined the nuclear localization of AGO1 upon transient depletion of *NEAT1* by antisense oligo (ASO) that targets both *NEAT1* isoforms. Strikingly, we observed a marked disappearance of AGO1 nuclear paraspeckle signals in *NEAT1* depleted cells (Fig. 7A). Next, by using CRISPR/Cas9 we generated *NEAT1* knocked-out (KO) human HAP1 (near-haploid) cell line (Fig. 6A). We used the HAP1 cell line because of the ease of genome editing in these cells. By combining *NEAT1* RNA FISH with SFPQ immunofluorescence, we confirmed the complete disappearance of paraspeckles in the selected *NEAT1-KO* HAP1 cell lines (Supplemental Fig. S10a). Consistent with *NEAT1* transient depletion, we observed a substantial adverse effect on AGO1 nuclear localization also in *NEAT1-KO* cells (Fig. 6B).

**Figure 6.**
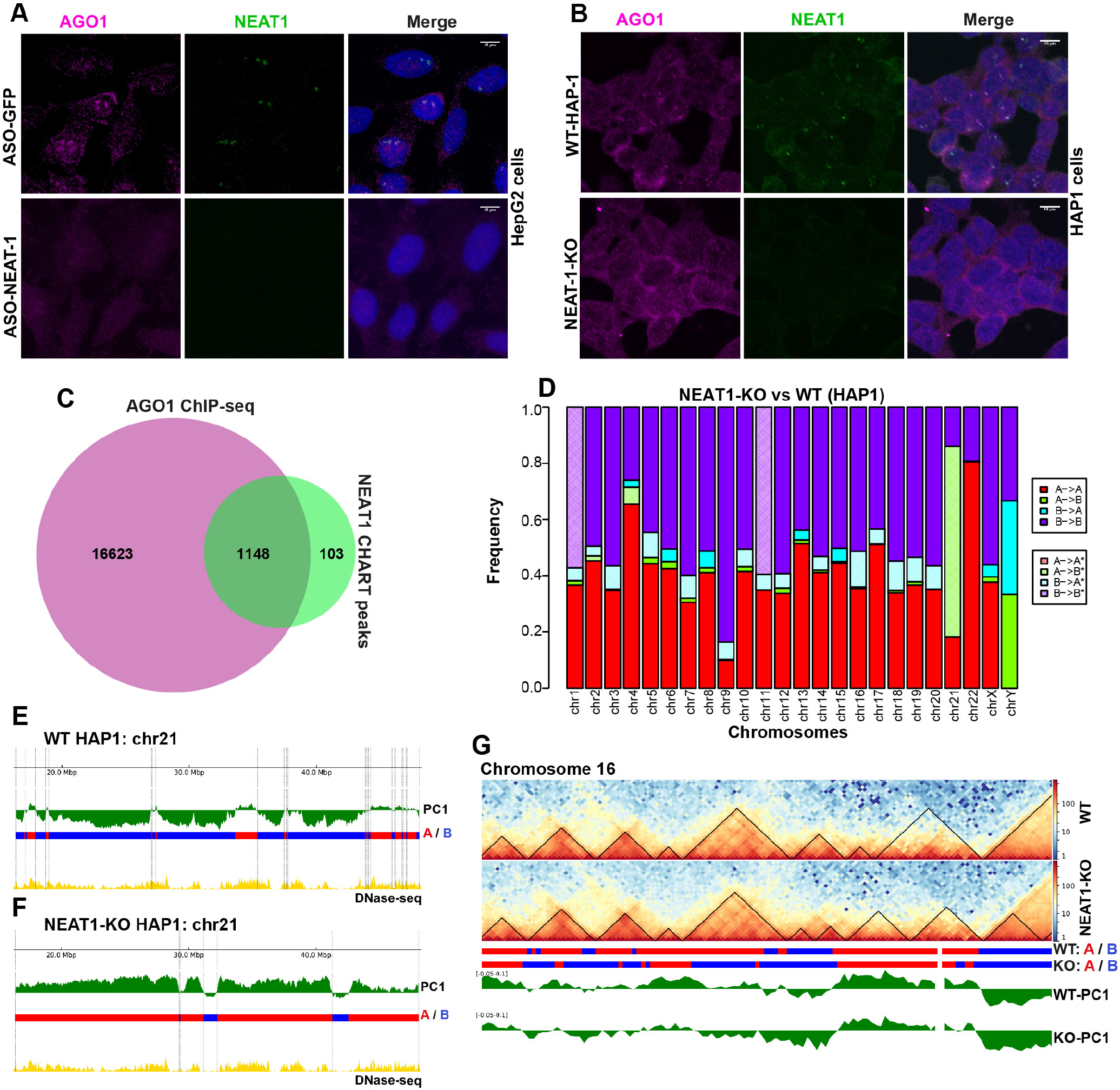
Loss of *NEAT1* affects AGO1 paraspeckle localization and higher-order chromatin organization. (*A*) Depletion of *NEAT1* by antisense oligo (ASO-NEAT1) affects AGO1 nuclear localization in HepG2 cells. GFP antisense oligo (ASO-GFP) was used as negative control. (*B*) Similar to (*A*) *NEAT1* knockout (KO) HAP1 cell lines displayed a diffuse AGO1 signals compared to WT-HAP1. (*C*) Venn-diagram shows overlapping of AGO1 ChIP-seq peaks with *NEAT1* CHART-seq data (Data accession# GSM1411207 and GSM1411208). (*D*) Bar chart showing A/B compartment changes between *NEAT1-KO* and WT HAP1 genomes. “A−>A” and “B−>B” mean no change in the compartment A and B respectively. “A−>B” represents compartments that are A-type in NEAT1-KO and B-type in WT HAP1. “B−>A” represents compartments that are B-type in NEAT1-KO and A-type in WT HAP1. Significant compartment changes are shown with same-color hatching (* Fisher test, FDR<0.05). (*E-F*) First eigenvector values of chromosome 21 used to define A/B chromatin compartments, positive values represent A-type (red) and negative values represent B-type (blue) compartments in WT-HAP1 (e) and *NEAT1* knockout HAP1 (*F*). Dotted lines delimit compartment borders. Active marks represent DNase-seq peaks (Data accession#: GSM2400413 and GSM2400414) in HAP1 cell. (*G*) Representation of TADs and compartments in Chromosome-16 (locus 50,000,000-60,000,000) identified in WT (top) and *NEAT1* knockout (bottom) HAP1 cells.

**Figure 7.**
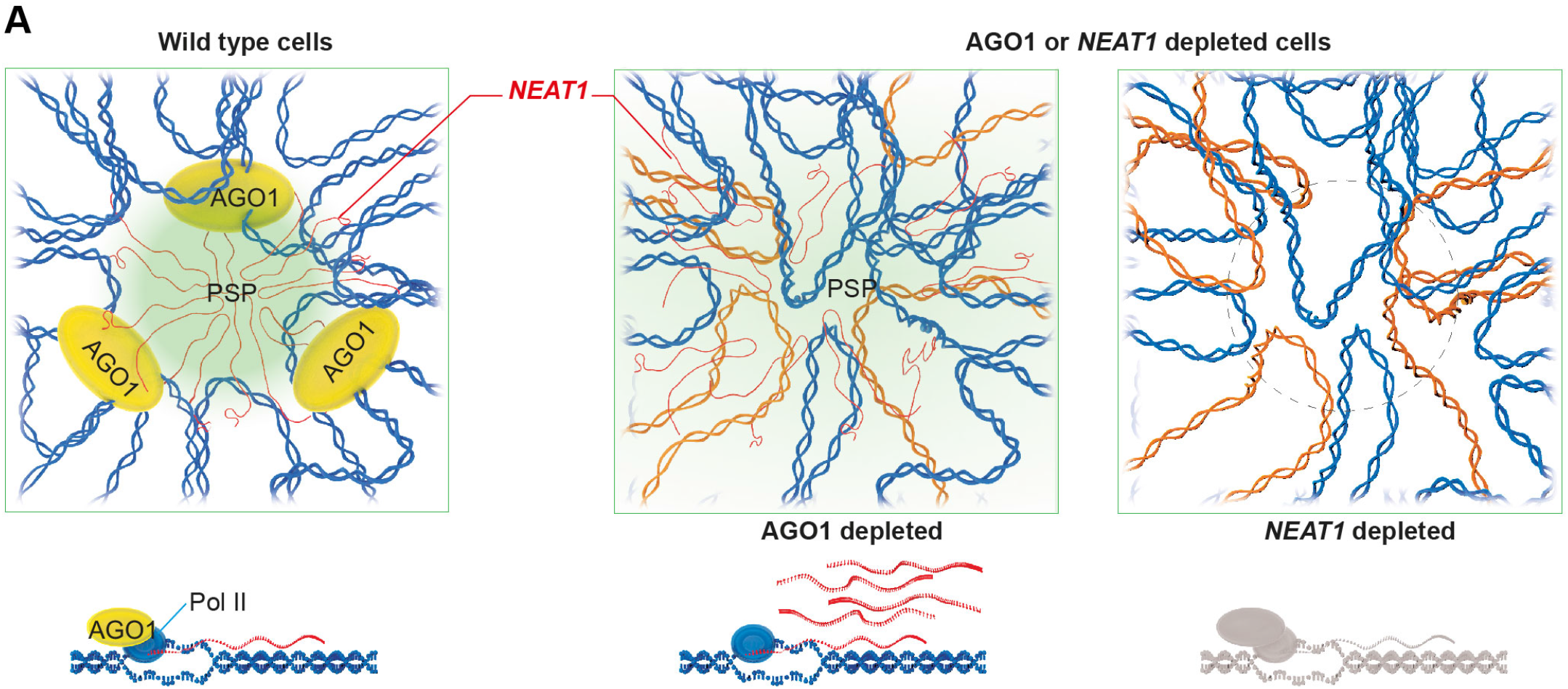
A model that shows AGO1 together with *NEAT1* lncRNA paraspeckle act as 3D genome organizers. (*A*) AGO1 in combination with *NEAT1* lncRNA and paraspeckle proteins (PSPs) facilitates trans-chromosomal interactions and establishes an epi-genomic environment that properly maintains TADs and compartments. AGO1 or *NEAT1* depleted cells show derangement in A/B compartments. AGO1 also binds to the promoter of *NEAT1* and controls its expression (lower panel). The depletion of AGO1 induces *NEAT1* expression and results in disorganized paraspeckles and *NEAT1* lncRNA (left). Similarly, *NEAT1* knockout cell lines display disorganized 3D genome structure (right).

*NEAT1* was reported to be enriched on active chromatin by CHART-seq in human cells (West et al. 2014). Our data show that AGO1 co-localizes with *NEAT1* across active sites and its depletion affects paraspeckle integrity and chromatin organization. We observed that around 94% of *NEAT1* CHART-seq peaks overlapped with AGO1 binding (ChIP-seq) at active sites (Fig. 6C). This finding suggests that the combinatorial action of *NEAT1* paraspeckles and nuclear AGO1 is required for correct genome architecture.

To test this hypothesis, we examined how the loss of NEAT1-paraspeckles affects chromatin organization using Hi-C in wild-type (WT) and *NEAT1-KO* HAP1 cell lines. We determined significant interaction matrices with the HiC-Pro pipeline (Servant et al. 2015) (Supplemental Fig. S10b-e), differential interactions by diffHiC (Lun and Smyth 2015), and (A/B) compartment analysis by principal components. The differential Hi-C analysis identified 3,567 bins showing a significant (FDR<0.05, logFC > 1.2) gain in intra-chromosomal interactions in *NEAT1-KO* cells.

Furthermore, comparing genome-wide compartments between *NEAT1-KO* and WT-HAP1 cells identified significant changes in A/B chromatin compartments (Fig. 7D). Notably, we observed switching of A to B compartments (Fisher’s test, FDR < 0.05) in the majority of the chromosomes (Fig. 6D). We also found B to A compartment transition across multiple regions, specifically on chromosome 2, 4, and 21 (Fig. 6D). As an example, the PC1 values illustrate the B to A compartment switching on chromosome 21 between WT and NEAT1-knockout HAP1 in Fig. 6E-F. Next, we checked the effect of NEAT1-knockout on TADs. Similar to what observed in AGO1 depleted cells, also in this case we detected “split or fused” topologically associated domains (TADs) configuration, as illustrated in one example region on chromosome 16 (Fig. 6G).

## Discussion

In this work, we report a link between nuclear AGO1, *NEAT1* lncRNA, paraspeckles formation, and higher-order chromatin topological structure. The role of RNA as an essential component of the nuclear architecture has been recognized for a long time (Holmes et al. 1972; Nickerson et al. 1989). Recent paradigmatic works showed Xist (Cerase et al. 2015) and Firre (Hacisuleyman et al. 2014) lncRNAs to mediate both cis- and trans-chromosomal interactions. Furthermore, CoT1 repeat-derived RNA fraction was shown to be essential for the integrity of chromatin organization (Hall et al. 2014). Our RIP-seq analysis identified several RNA moieties that are associated with nuclear AGO1, including lncRNAs, mRNAs and short RNAs. While we do not exclude binding to short RNAs, because of the association of AGO1 with active regulatory cis-elements our work focused on other RNA moieties and in particular lncRNA, as a potential bridge between AGO1, chromatin association, and gene expression. Among all, *NEAT1* lncRNA resulted to be the most represented one. *NEAT1* lncRNA is the key organizer of paraspeckles, containing various RNAs and RNA binding proteins (RBPs) (Prasanth et al. 2005; Hirose et al. 2014; Imamura et al. 2014). Indeed the data presented revealed: I. AGO1 co-localization with *NEAT1* containing paraspeckles, II. AGO1 association with essential paraspeckle components, III. AGO1 *NEAT1* direct binding (ChIRP, CLIP), IV. impact on paraspeckle integrity upon AGO1 knockdown V. loss of AGO1 paraspeckles localization upon *NEAT1* depletion.

Along with *NEAT1*, we also identified *MALAT1* lncRNA in our RIP-seq and CLIP analysis. Also, AGO1 binds to the promoter of both *NEAT1* and *MALAT1 genes* that are located adjacent to one another on chromosome11. *MALAT1* and *NEAT1* lncRNAs localize to distinct nuclear bodies the nuclear speckles and paraspeckles respectively. These nuclear bodies often localized adjacent to one another in the nucleus (Fox et al. 2002) and also share essential protein components, like PSF and NONO (Saitoh et al. 2004). The *NEAT1* lncRNA has also been shown to localize to nuclear speckles upon inhibition of transcription elongation (Sunwoo et al. 2009). The significant number of genomic loci that are co-enriched by both *NEAT1* and *MALAT1* (West et al. 2014) also overlapped with AGO1 binding sites. These observations suggest a significant overlap between *NEAT1, MALAT1*, and AGO1 nuclear function. Further work should investigate the cross-talk between speckle and paraspeckle and its impact on 3D genome organization in AGO1 depleted cells.

The mechanistic explanation of the impact of AGO1 depletion on nuclear and higher order chromatin organization remains to be understood. Our findings show that AGO1 interacts with both *NEAT1* TSS and *NEAT1* lncRNA and its depletion induces *NEAT1* over-expression but adversely affect paraspeckle formation indicates that AGO1 impacts both *NEAT1* expression and paraspeckles assembly. The disruption of genomic compartments and TAD boundaries may lead to deregulated gene expression (Nora et al. 2012; Phillips-Cremins et al. 2013; Seitan et al. 2013; Barutcu et al. 2015). AGO1-knockdown CAGE-seq analysis shows that the majority of gene expression changes correlate with perturbed topologically interacting sites and alterations in TAD structure. The differential expression of various non-coding RNAs (Isoda et al. 2017) including *NEAT1* may contribute to topological organization. Consistently, the effect of AGO1 depletion, Hi-C analysis in *NEAT1-KO* cells reveals similar changes in higher-order chromatin architecture. A recent report showed a link between phase-separation and NEAT1-dependent paraspeckles assembly (Yamazaki et al. 2018). Thus, it is tempting to speculate that AGO1-NEAT1 lncRNA interaction might regulate the differential aggregation states of specific cis-regulatory elements with impact on transcriptional output.

Regarding other AGO proteins, currently, it is unclear whether different AGO proteins (AGO1-4) might play a similar function in the nucleus. AGO1 depletion appears to increase AGO2 nuclear levels and also to affect the nuclear redistribution of other RNAi factors (Matsui et al. 2015). Our data show that compared to AGO2-3, AGO1 proteins mostly enrich in the chromatin fractions. Moreover, unlike AGO1, AGO2 ChIP-seq analysis does not detect widespread genomic distribution (Huang et al. 2013). While we cannot exclude any technical problems related to the ChIP efficiency of AGO2 antibody, the difference in bulk chromatin binding may reflect AGO2 peripheral nuclear localization (Huang et al. 2013). Notably, a recent study (van Eijl et al. 2017) reported about the cross-reactivity of another AGO2 antibody (11A9) with SMARCC1, which was previously used for ChIP (Ameyar-Zazoua et al. 2012) and protein immunoprecipitation (Carissimi et al. 2015). Therefore, the nuclear function and genome-wide association of AGO2 remains controversial. However, direct or indirect mechanistic evidence that AGO2/3 would compensate for AGO1 nuclear function remain to be reported. Additionally, our data show that the essential paraspeckle proteins specifically associate with AGO1 but not AGO2 nuclear complexes, indicating different functions for AGO1 and AGO2 in the nucleus. Importantly, in contrast to AGO2 (Meister et al. 2004) and AGO3 (Park et al. 2017), human AGO1 does not contain a functional slicing domain (Wu et al. 2008). Thus its nuclear function appears to be independent of RNA-processing, but instead, it depends on RNA binding.

Finally, our data demonstrate that depletion of AGO1 or *NEAT1* leads to convergent phenotypic consequences namely widespread alterations of fundamental higher-order genomic compartmentalization and TADs organization (Fig. 7). Notably, there is a strong link between overexpression of NEAT1-lncRNA and several aggressive types of cancer (Chakravarty et al. 2014; Chen et al. 2015; Ma et al. 2016) that also display dramatic alterations in 3D genome organization (Barutcu et al. 2015; Achinger-Kawecka et al. 2016; Taberlay et al. 2016). The reported connection with AGO1 provides a novel insight towards understanding NEAT1-dependent mechanism operating in the development of this disease possibly via nuclear architecture.

## Methods

### Cell culture

HepG2 and HEK-293 cells were cultured in EMEM medium (Sigma, Cat # M0643) supplemented with 10% FBS (Invitrogen, Cat # 26140-079), and HAP1 cells were grown in Iscove’s Modified Dulbecco’s Medium (IMDM, GIBCO, Cat.No. 12440-053) with 10% FBS. 1% penicillin and streptomycin (Euroclone, Cat # ECB3001D) was added to culture media. Cell cultures were maintained at 37°C and 5% CO2 (see supplemental methods for detail).

### Fluorescent *in situ* hybridization (RNA-FISH) and Immunocytochemistry

The probes for *NEAT1* lncRNA were synthesized by *in-vitro* transcription of antisense RNA using linearized plasmids containing a *NEAT1* fragment (+1 to +1,000). The experiments were performed as described previously (Naganuma et al. 2012; Kawaguchi et al. 2015).

### Chromatin fractionation

Cells were lysed in RIPA buffer (10 mM Tris pH 8.0, 1 mM EDTA, 0.5 mM EGTA, 1% Triton X-100, 0.1% Sodium deoxycholate, 0.1% SDS, 150 mM EDTA and 0.5 mM PMSF, 10 U/ml Superase-In, 1X Protease inhibitor cocktail), incubated for 30 minutes on ice, and spun at 13000 rpm for 10 min to collect supernatant as total fraction (T). For chromatin fractionation, cells were lysed in CSKI buffer (10 mM Pipes pH 6.8, 100 mM NaCl, 1 mM EDTA, 300 mM Sucrose, 1 mM MgCl_2_, 1 mM DTT, 0.5% Triton X-100, 10 U/ml Superase-In, 1X Protease inhibitor cocktail) for 15 min on ice (vortexed occasionally) and spun at 3000 rpm for 5 min to remove the soluble fraction (S). Pellet was resuspended in CSK II buffer (10 mM Pipes pH 6.8, 50 mM NaCl, 300 mM Sucrose, 6 mM MgCl_2_, 1mM DTT), treated with DNase I (Promega, M610A) for 25 min at 30°C, and followed by extraction with 250 mM (NH_4_)_2_SO_4_ for 10 min. The extract was centrifuged at 1200g for 10 min at 4°C to collect the supernatant as chromatin bound fraction (CB).

### nanoCAGE-seq

After cellular fractionation of HepG2 cells as described above, Trizol was added to the total and chromatin bound fractions for RNA extraction. Samples were processed for nanoCAGEseq as described previously (Salimullah et al. 2011).

### ChIP-seq

HepG2 cells were cross-linked with 2% formaldehyde on shaking platform for 12 min at room temperature. The reaction was quenched with 125 mM of glycine and cells were collected by scraping. The pellet was further processed by incubating in lysis buffer (5 mM HEPES/KOH pH 8.0, 85 mM KCl, 0.5% NP-40, 0.5 mM PMSF) for 30 min on ice. Pelleted nuclei were resuspended in shearing buffer (50 mM Tris/HCl pH 8.0, 1 mM EDTA, 0.1% SDS, 0.5% DOC, 0.5 mM PMSF), passed through 1 ml syringe for 10-15 times, and sonicated with Diagenode Bioruptor (total 3 cycles for 7 min each with settings 30 sec ON and 30 sec OFF at maximum amplitude). To remove debris, sonicated samples were spun at 16000xg for 20 min at 4°C. Supernatant was collected and chromatin DNA fragment size (average 500 bp) was checked after decrosslinking at 65°C for 4 hrs on 2% agarose gel. Chromatin suspension was mixed with 1X IP buffer (1 mM Tris/HCl pH 7.4, 0.1 mM EDTA, 0.1% Triton X-100, 0.05% DOC and 150 mM NaCl, 0.5 mM PMSF), AGO1 antibody (Wako, Clone 2A7) and rotated overnight on a rocker wheel at 4°C. Immuno-complexes were captured with Dyna beads Protein G (Invitrogen) for 2 hrs on a rocker wheel at 4°C. Further, beads were washed with 2X Low Salt buffer (20 mM Tris/HCl pH 8.0, 2 mM EDTA, 0.1% SDS, 1% Triton X-100, 150 mM NaCl, 0.5 mM PMSF), 2X High Salt buffer (20 mM Tris/HCl pH 8.0, 2 mM EDTA, 0.1% SDS, 1% Triton X-100, 500 mM NaCl, 0.5 mM PMSF), 1X LiCl buffer (10 mM Tris/HCl pH8.0, 10 mM EDTA, 1% NP-40, 250 mM LiCl, 0.5 mM PMSF), and 1X TE (1 mM Tris/HCl pH 8.0, 1 mM EDTA). Finally, immunocomplexes were eluted (1 mM Tris/HCl pH 8.0, 10 mM EDTA, 1% SDS) at 65°C for 20 min, cross-links were reversed at 65°C overnight, treated with RNase A, proteinase K and then DNA was extracted with Phenol:Chloroform:Isoamyl alcohol (25:24:1). The extracted ChIP-DNA was processed for high-throughput library preparation.

### AGO1 RIP from chromatin fraction

The AGO1 RIP-seq was performed following the above ChIP experiment until the last washing step as described in the protocol. The whole procedure was carried out at 4°C and all the buffers were prepared in DEPC (0.1 % (v/v) and supplemented with Superase-In (Ambion – AM2696) 10U/ml. After proteinase K treatment (0.4 mg/ml) on beads at 37°C for 30 min, the samples were processed for the reverse cross-linking at 65°C for 2 hrs. Finally, the RNAs were extracted using TRI reagent and processed for library preparation.

For UV cross-linking IP combined with qRT-PCR, cells were cross-linked on ice with 254 nM UV-C at 0.3 J/cm2, and the rest of the experiment was performed as described previously (Sasaki et al. 2009).

### Preparation of Hi-C libraries and generation of contact matrices

Hi-C experiments were carried out as previously described (Lieberman-Aiden et al. 2009; Belton et al. 2012), with minor modifications(Schoenfelder et al. 2015). Briefly, 25 million cells were fixed with 2% formaldehyde and lysed. Following HindIII digestion and biotinylation of DNA ends, ligation was performed inside the nuclei. After de-crosslinking, the DNA was purified, sheared, and pull-down performed with streptavidin beads. Hi-C libraries were prepared for sequencing on HiSeq 4000 (Illumina). After sequencing, Fastq data were processed through HiC-Pro pipeline(Servant et al. 2015) for generating contact matrices at different resolutions. We obtained a total of around 190 million valid interaction pairs, ~95 million for siCtrl and ~95 million for siAGO1 HepG2 cells (combined replicates) (Supplemental Fig. S6a-d), the two replicates in each condition showed high degree of similarity and reproducibility (Supplemental Fig. S6e). Similarly, we obtained around 466 million valid pairs, ~200 million for WT-HAP1 and ~266 million for NEAT1-KO HAP1 cell lines (Supplemental Fig. S10b-c).

### Bioinformatics analyses

The detail bioinformatics analyses are provided in the supplemental methods.

## Data access

The high-throughput sequencing data including CAGE-seq, Small RNA-seq, ChIP-seq, RIP-seq, and Hi-C-seq have been deposited at the Sequence Read Archive (SRA, http://www.ncbi.nlm.nih.gov/sra/), which is hosted at the NCBI, under the SRA accession number SRP115598 (BioProject ID# PRJNA398595).

## Acknowledgments

We are grateful to Hakan Ozadam and Johan Gibcus from Job Dekker group (University of Massachusetts Medical School, USA) for initial help in Hi-C analysis; Riccardo Aiese Cigliano (Sequentia Biotech) for help in bioinformatic analysis; Ana Maria Suzuki (RIKEN) for help in CAGE-seq; Heno Hwang (scientific illustrator) at KAUST for his help in drawing the image (Figure-8); Christian Froekjaer Jensen for critical reading of the manuscript; KAUST Bioscience Core Lab for providing sequencing facility. The work was supported by EPIGEN-CNR (Italian Ministry of University and Research) and King Abdullah University of Science and Technology (KAUST) to V.O. The grants from the MEXT of Japan to TH (26113002 and 17H03630). The grant for Joint Research Program of IGM, Hokkaido University.

## Author contributions

M.S conceived this study, designed and performed experiments, analyzed the data and wrote the manuscript with input from all the authors. K.M.P designed and performed experiments. S.A.A contributed in Hi-C experiments. A.F. contributed in RIP-seq experiments. M.T and H.K. performed computational analyses of RIP-seq, CAGE-seq and ChIP-seq data. Y.G and L.S performed computational analyses of RIP-seq, ChIP-seq and CAGE-seq data. T.Y provided the HAP1 cells (NEAT1 knockout and wild type). T.M and T.Y contributed in the RNA FISH experiments. B.F helped in western blotting. T.H conceived and designed the experiments related to paraspeckles. P.C and H.K. produced and analyzed the CAGE-seq experiment. V.O. conceived this study, designed experiments, and wrote the manuscript.

