## Supplemental Data for "AGO1 in association with *NEAT1* lncRNA contributes to nuclear and 3D chromatin architecture in human cells"

#### Supplemental Results

##### AGO1 nuclear localization and its knockdown by siRNAs in HepG2 cells

In order to understand the function of endogenous nuclear AGO proteins at chromatin level, we first examined the association of AGO proteins (AGO1-4) with chromatin in HepG2 cells, by western blot and immunofluorescence analysis. After cellular fractionation into total (T), soluble (S), chromatin (CB) and matrix bound (CR) (Supplemental Fig. S1a), western-blot analysis revealed that AGO1 was more enriched compared to AGO2, AGO3 and AGO4 in the chromatin-bound fraction (Supplemental Fig. S1b). Further analysis of AGO1 cellular distribution by immunofluorescence (IF) confirmed AGO1 presence spread across the nucleus (Supplemental Fig. S1c). These results indicate that endogenous nuclear AGO1 has chromatin related function in human cells.

To investigate role of AGO1 in genome organization and gene expression regulation, we knockdown (KD) AGO1 with a pool of four siRNAs (siAGO1) and performed a control knockdown using a non-specific scramble siRNA (siCtrl) in HepG2 cells. The efficiency of knockdown for each individual single siRNA was also tested. AGO1 depleted cells showing a similar pattern of cell viability (Supplementary Fig. S1d) and cell cycle profiles compared to control KD cells (Supplementary Fig. 1e) were processed for further analysis We confirmed the down-regulation of AGO1 by western blot, qRT-PCR and immunofluorescence analysis (Supplemental Fig. S1e-i). Next to check the specificity of AGO1 siRNAs pool, we performed western blot analysis for AGO1, AGO2 and AGO3 after transfection. We did not find a decrease in the total level of AGO2 and AGO3 upon AGO1 depletion by siRNAs (Supplemental Fig. S1j).

**AGO1 associates with active genomic regulatory landscape and its knockdown alters transcriptional output**

To identify genome-wide transcriptional changes upon AGO1 depletion, we performed CAGE-seq (total and chromatin bound RNAs) using three replicates for WT, siCtrl and siAGO1 HepG2 cells. All three replicates in each condition showed a similar pattern as represented in PCA plot (Supplemental Fig. S2a). By comparing CAGE-seq data from total RNA and chromatin-bound RNA, we observed 171 common differentially expressed genes in the two fractions (Supplemental Fig. S2b). Considering only the highly significant perturbed genes (FDR ≤ 0.05), we found that 512 genes were up regulated, whereas 226 genes showed down-regulation in total RNA-CAGE. Similarly, 213 up-regulated genes and 170 down-regulated genes were found in CAGE data from chromatin bound RNAs (Supplemental Fig. S2c).

To identify AGO1 genomic distribution and to determine whether AGO1 directly regulates the perturbed promoters observed upon AGO1-depletion by CAGE-seq, we performed ChIP-seq analysis. AGO1 ChIP-seq data analysis revealed approximately 80% mapping rate, and we identified 17,771 reproducible AGO1-specific peaks. Examining the distribution of these peaks, we found that AGO1 was highly enriched in many active regions of the genome, such as promoters, enhancers, and also other genic regions (i.e. introns, exons, and UTRs) (Supplemental Fig. S3c-d).

#### Supplemental Methods

##### Antibodies

Mouse monoclonal anti-hDICER1 (12B5/4C6), anti-AGO1, anti-AGO-2, anti-AGO3, and anti-AGO4 sup (a kind gift from Mikiko Siomi), anti-AGO1 (clone 2A7, Wako), anti-SFPQ (RN014MW, Ribonomics), anti-HNRNPK (SAB4501424, Sigma), and anti-TLS/FUS (ab23439, Abcam), and anti-PSPC1 (ab104238, Abcam), β-Tubulin antibody (#2146, Cell Signaling), and Histone H3 antibody (#9715, Cell Signaling), goat anti-NONO /p54NRB (EB07246) were used as primary antibody for western blotting and or immunofluorescence. Alexa flour 647 Goat anti-mouse IgG (H+L) (ab150115, Abcam) and Alexa flour 488 Donkey anti-mouse IgG (H+L) (Molecular probe, A21202) were used as secondary antibody for immunofluorescence. Mouse monoclonal anti-AGO1 (clone 2A7, Wako) for ChIP experiment and control validation by western blot and immunofluorescence (Supplemental Fig. S1e-g).

**Transfection and flow cytometry**

AGO1 knockdown experiments were performed using a pool of four siRNAs (FlexiTube GeneSolution GS26523, the individual catalogue numbers for each siRNA are SI00377454, SI00377447, SI00377440, and SI00377433) against AGO1. The efficiency of AGO1 down-regulation and its impact on gene expression was also tested for each single siRNA (Supplemental Fig. S1k-m). For transfection HiPerfect reagent (Qiagen) was used according to the supplier's protocol. After 72hrs of incubation, cells were harvested for further analysis. Cell viability was assessed using alamarBlue cell viability reagent (Cat# DAL1025) according to the manufacture's instructions. For flow cytometry, cells were pelleted and fixed in ice cold 70% ethanol. After fixation the cell pellet was resuspended in the PI staining solution (0.1% Triton X-100, 10 µg/mL propidium iodide and 100 µg/mL RNase A). Cell-cycle analysis was performed using BD FACSCanto II system and experiments were repeated three times.

The phosphothioate-converted antisense oligonucleotides (ASO) against *NEAT1* or negative control GFP were used as describe previously (Sasaki et al. 2009). The transfection into HepG2 cell nuclei was performed by nucleofection according to standard protocol (Hirose and Mannen 2015).

**Establishment of *NEAT1* knockout HAP1 cell lines by CRISPR/Cas9**

HAP1 cell lines were purchased from Horizon Discovery. *NEAT1* knockout cell lines were produced by CRISPR mediated deletion of the complete *NEAT1* gene locus using two sgRNAs that target the 5'- and 3'-ends of *NEAT1.* The guide RNAs were designed using CRISPR Design website (crispr.mit.edu). PX330-derived vector (PX330 from Addgene) harboring these two sgRNA and Cas9 sequences (2 mg) were co-transfected with pcDNA6/TR plasmids (0.2 mg) containing blasticidin resistant gene (Thermo Fisher Scientific) into HAP1 cells (1.5 x $10^5$ cells) by Nucleofector Kit V (Lonza) with a Nucleofector device (Lonza) using a program "X-005" according to the manufacturer's instruction. To enrich the plasmid-transfected cells, the HAP1 cells were treated with 20mg/ml blasticidin (InvivoGen) for 3 days, starting one day after transfection. Subsequently, the cells were diluted into 96-well plates for selection of single clones. The genomic DNA from the selected clones was extracted after lysing the cells by proteinase K treatment (200 mg/ml proteinase K [Roche], 20 mM Tris-HCl, pH 8.0, 5 mM EDTA, 400 mM NaCl, 0.3 % SDS at 55°C for 1 hour, followed by proteinase K inactivation; 95°C for 1 minutes). Then the extracted DNA was subjected to PCR analysis to amplify the genomic regions flanking the guide RNA target sites for detecting deletions by using KOD FX Neo enzyme (TOYOBO). Positive clones were further confirmed by sequencing.

**Immunofluorescence**

For AGO1 staining, HepG2 cells were grown on cover slips and fixed with 4% formaldeyhyde. Cells were permeabilized with 0.4% Triton-X100, blocked and incubated with the primary antibody (1:400 dilution) over night at 4°C. Next day, cells were stained with secondary antibody at room temperature for one hour. DAPI (Invitrogen) was used for nuclear staining. Images were acquired using confocal microscope (Zeiss).

**Cellular fractionation and western blot**

Cells were lysed in hypotonic buffer (10mM Tris pH8, 10mM KCl, 1mM EDTA, 1mM EGTA, 1% NP-40, 10% glycerol, 0.5mM PMSF) for 30 minutes on ice and spun at 4000rpm for 10min to collect supernatant as cytoplasmic fraction. Pellet was washed twice in cytoplasmic buffer, and then resuspended in sucrose buffer (20 Tris (pH 7.65), 15mM KCl, 60mM NaCl, 0.34M Sucrose, 0.15mM spermine, 0.5 mM spermidine). High salt buffer (20mM Tris (pH 7.65), 25% glycerol, 1.5 mM MgCl2,, 0.2mM EDTA, 900mM NaCl) was added drop by drop while vortexing to a final concentration of 0.3M and incubated on wheel for 30 minutes at 4°C. Lysed nuclei were centrifuged at 4000rpm for 10min at 4°C to remove the soluble nuclear fraction. Mnase digestion was performed using Micrococcol nuclease (0.5U/μl sigma) 5μl per gram of cells at 37°C for 10 minutes and then sonicated and centrifuged at 35000rpm for 1hr at 4°C to collect the chromatin fraction. Protein samples from all fractions were denatured in 1X Laemmli buffer (100mM DTT) and separated on 4-12% gradient Bis-Tris gels (NuPAGE), and then transferred onto a nitrocellulose membrane (GE Healthcare) for 2 hours. The membrane was blocked with 5% milk in PBST, incubated in primary antibody overnight, washed with PBST, and incubated with secondary antibody for 1 hour. Western-blot signals were visualized using ECL Reagent (Amersham) and ChemiDoc Imaging System (Bio-Rad).

**RT-qPCR**

RNA was extracted with TRI reagent (Sigma, Cat # T9424) and reverse transcribed using the QuantiTect reverse transcription kit (Qiagen, Cat # 205311). cDNA were then amplified by quantitative PCR on CFX96 Real-Time PCR Machine (Bio Rad) using the SYBR Select Mastermix (Applied Biosystems). For relative quantification of total RNAs, the geometric mean of 18S rRNA, GAPDH, and Actin-B mRNA levels were used for normalization. Human AGO1 specific primer was purchased from Qiagen (Cat#

QT00006370) and reference genes (18S, GAPDH, and ActB) primers were used from geNorm Syber green kit (Primerdesign Cat# ge-SY-12).

**Tandem affinity purification and mass spectrometry**

The soluble nuclear extracts and chromatin bound extracts were prepared from stable HEK-293 cell lines expressing either AGO1 or AGO2 proteins fused to C-terminal FLAG and HA epitope tags (tagged protein are denoted as e-AGO1 and e-AGO2). The e-AGO1 and e-AGO2 protein complexes were purified by double-immunoaffinity purification procedure from nuclear soluble and chromatin fraction with anti-Flag M2 antibody-conjugated agarose (Sigma), followed by anti-HA purification. Interacting protein partners of purified complexes (AGO1-com and AGO2-com) were identified by mass spectrometer (Bioscience Core laboratory, KAUST). The interactions with specific partners were further confirmed by western blotting.

***NEAT1* ChIRP and western blotting**

*NEAT1* ChIRP followed by protein isolation was performed as described previously (Chu et al. 2015). Briefly, 500 million cells (per ChIRP reaction) were cross-linked with formaldehyde, target RNA was retrieved with Magna ChIRP *NEAT1* probes set (even, odd) (Cat. # 03-308) or Magna ChIRP negative control probe (LacZ, part # CS216572), and proteins were eluted with biotin. The enriched proteins were used for silver staining and western blotting.

**Small RNA-seq**

The small RNA-seq libraries were generated from total RNA with three replicates in each condition (siCtrl and siAGO1 HepG2) using Illumina TruSeq small RNA protocol. These libraries were sequenced on Illumina HiSeq2000. After QC and trimming of low quality bases and adapters, the fastq files were uploaded to the online small RNA-seq analysis tool Oasis 2.0 (https://oasis.dzne.de) (Rahman et al. 2018) for detection, differential expression, and classification of small RNAs. We identified a list of significant (p-value 0.01) differentially expressed miRNAs.

For integration with CAGE-seq data, we used the mirTarBase tool (mirtarbase.mbc.nctu.edu.tw/php/) (Chou et al. 2018) to identify the targets of differentially expressed miRNAs in the list of differentially expressed genes from AGO1 knockdown CAGE-seq data.

**Bioinformatics analyses**

**CAGE-seq data analysis**

The CAGE libraries were generated using RNA extracted either from chromatin or total fraction with three replicates in each condition (siCtrl and siAGO1 HepG2). These libraries were sequenced using HiSeq 2000 with single read of 50 bp lengths. The low quality bases and adapter were trimmed using cutadapt tool. The clean reads were then mapped on to the human reference genome hg19 from UCSC. To get the tag counts that mapped onto the genomic loci, the mapped bam files were analysed using the R package CAGEr (https://bioconductor.org/packages/release/bioc/html/CAGEr.html). Expression tables were generated by counting the number of mapped reads within 500 bases of an UCSC (http://hgdownload.cse.ucsc.edu/goldenPath/hg19) annotated 5 prime ends as expression from the transcript in CAGE. The number of tags for each sample was used as input counts and uploaded to the DESeq2 for gene expression calculation (https://bioconductor.org/packages/release/bioc/html/DESeq2.html). The up- and down- regulated genes were tabulated with significance FDR threshold of 0.05.

ChIP-seq data analysis

After sequencing ChIP-seq reads of quality score less than 20 and length of 30 bp were removed (http://www.citeulike.org/user/mvermaat/article/13260426) using sickle. We aligned the reads to the human genome assembly GRCh37 (hg19) using Bowtie(Langmead and Salzberg 2012) (version 2.2.5) using default parameters, and results were converted to sorted bam using SamTools(Li et al. 2009) (version 1.2). HMCan peak calling tool (Ashoor et al. 2013) was used for ChIP-seq peak calling. This tool is used for cancer cells to normalize reads for GC and copy number variation. Peak calling was then followed by irreproducibility discovery rate (idr) analysis (Li et al. 2011). For peaks annotation, we used GENCODE (Harrow et al. 2012) (version 19) annotation of the genome. ChIP-seq peaks were annotated using Bedtools (Quinlan and Hall 2010) and BEDOPS (Neph et al. 2012). Gene expression was measured as the sum of normalized CAGE tags within 500 bp from TSS. For integration with ChIP-seq, CAGE signals around ChIP-seq peaks were computed using bwtools (Pohl and Beato 2014) (version 2.17).

**RIP-seq analysis**

The RIP-seq libraries were prepared from two replicates of chromatin-bound AGO1 associated RNAs and mock sample RNAs (control) for 100 and 400 fragment sizes. Raw sequencing reads were aligned to the UCSC human genome release hg19 using tophat2 (v2.0.14) (Kim et al. 2013). Also, a reference GTF obtained from Illumina iGenomes was used and supplied to tophat2 (https://support.illumina.com/sequencing/sequencing_software/igenome.html). In order to identify enriched peaks, the target and mock reads for each fragment size and for each replicate independently were first aligned. They were then used as input for Piranha(Uren et al. 2012) (v1.2.1) with the following command line options "-s -l". Only consistent peaks, identified through genomic overlap of at least 1 base were retained using BEDTools. Figures were generated using R and ggplot2.

#### Hi-C data analysis

##### Differential interaction analysis

Genome-wide Hi-C matrices were loaded to the R package diffHiC (Lun and Smyth 2015) (version 1.6.0) to identify differential interactions between siAGO1 and siCtrl HepG2 cells and similarly between *NEAT1*-KO and WT HAP1 cell lines. Those interacting pairs of bins with an average read count less than 5 (considering all replicates) were filtered out. The LOESS method was used to normalise counts between libraries. The interacting pair of bins with a FDR value smaller than 0.05 were considered significant.

##### Identification of Topologically Associated Domains (TADs)

Columns with low number of counts in the genome-wide Hi-C matrices at 100kbp resolution were filtered out using TADbit (version 0.2.0.23) setting the parameter “min_count” to 10. Since TADbit fits the column count distribution into a polynomial distribution, columns with a number of counts smaller than the first antimode of the distribution, which cannot be smaller than the “min_count” parameter, are filtered out. Then, 100 kbp genome-wide matrices were normalized by visibility (1 iteration of ICE). Afterwards, the breakpoint detection algorithm of TADbit that returns the optimal segmentation of the chromosome under BIC-penalized likelihood was applied in order to identify Topologically Associating Domains (TADs). Hi-C matrices at 40 kbp resolution were processed per chromosome instead of genome-wide.

##### Identification of compartments

For compartment profiling, first columns with low number of counts in the genome-wide Hi-C matrices at 100 kbp resolution were filtered out using TADbit (version 0.2.0.23) setting the parameter “min_count” to 10. Then, 100 kb or 80kb (for HAP1) genome-wide matrices were normalized by the expected interactions at a given distance and by visibility (1 iteration of ICE). The correlation analysis was performed with TADbit tool to obtain the first eigenvector. In-house scripts computed A/B compartments from the first eigenvector, using 0 as threshold to differentiate both compartments and genes with a basemean expression higher than 20 as active marks to label compartments. Hi-C matrices at 40 kbp resolution were processed per chromosome instead of genome-wide.

**Heatmaps and multi-track plots**

The raw and normalized corrected matrices were plotted using hicPlotMatrix of HiCExplorer (ref) (version 1.3) (http://hicexplorer.readthedocs.io/en/latest/index.html). Corrected matrices underwent a low-count bin filtering and ICE normalization (500 iterations) prior to plotting. Multi-track plots were generated using HiCExplorer hicPlotTADs.

**TAD compartment switching**

TADs identified in siCtrl cells were intersected using the bedtools intersect with the first eigenvector of siCtrl and siAGO1 cells. Positive eigenvector values were attributed to A compartments while negative eigenvector values were attributed to B compartments. For each TAD and condition, the mean of the values of the corresponding eigenvector was calculated. TADs having a change in the sign of the calculated means were designated as TADs that switched compartments.

#### Supplemental Figure Legends

**Supplemental Figure 1 AGO1 subcellular localization and siRNA-mediated knockdown in HepG2 cells.** (a) Schematic representation of HepG2 cellular fractionation. (b) HepG2 cellular fractionation and western blotting using specific antibodies to detect AGO1, AGO2, AGO3 and AGO4 in different subcellular fractions (T = Total, S = Soluble, CB = Chromatin bound and CR = Chromatin released fractions or matrix associated). Tubulin and H3 antibodies were used as cytoplasmic and chromatin markers, respectively. (c) Detection of AGO1 nuclear localization in HepG2 cells, by immunofluorescence analysis. AGO1 specific antibody detects both chromatin associated and cytoplasmic endogenous AGO1. DAPI was used to stained nuclei (blue). Representative image is shown (scale bar 10µm). (d) Checking localization of AGO2 relative to paraspeckles in human HepG2 cells. AGO2 was detected by immunofluorescence and *NEAT1* RNA-FISH was used to visualize paraspeckles. (e-i) HepG2 cells were transfected with negative control siRNA (siCtrl) and siRNAs against

AGO1 (siAGO1) at 10nM concentration for 72 hrs. (e) Cell viability quantification by Alamar Blue (AB) assay for non-transfected and transfected cells (siCtrl and siAGO1). Viability is expressed as a percent of the control (non-transfected). For each condition, n = 3 different experiments (each with triplicates). (f) Cells were analyzed by flow cytometry after PI staining to measure DNA content. Flow cytometry analysis display similar cell cycle profiles for non-transfected, negative control transfected (siCtrl), and AGO1 siRNA transfected (siAGO1) cells. (g) Western blot showing AGO1 protein levels in siCtrl and siAGO1 cells, from two independent experiments. Tubulin was used as a loading control. (h) Bar graph shows the decrease in the level of AGO1 mRNA in siAGO1-transfected cells, as measured by RT-qPCR. Quantification of AGO1 mRNA level was performed with respect to three control RNAs (GAPDH, Actin-B, and 18s). (i) AGO1 signal disappearance upon knockdown. (j) Western blot showing AGO1, AGO2, and AGO3 protein levels in siCtrl and siAGO1 cells, from two independent experiments. Tubulin was used as a loading control. (k) Bar graph shows the reduction in the level of AGO1 mRNA upon treatment with single siRNA against AGO1, as measured by RT-qPCR. GAPDH and 18s were used as control RNAs. (l) Western blot showing decrease in protein level of AGO1 in cells transfected with individual siRNA. Tubulin was used as a loading control. (m) Changes in expression of selected target genes for each single siRNA treated cells were confirmed by qRT-PCR. Values were normalized relative to the geometric mean of 18S rRNA and GAPDH mRNA expression levels. Error bars indicate mean ± s.e.m from 2 independent experiments.

**Supplemental Figure 2 CAGE-seq and small RNA-seq integrative analysis in AGO1 knockdown cells.** (a) PCA (Principal Component Analysis) plot of CAGE-seq data showing the clustering of replicates for siCtrl, siAGO1 and non-transfected wild type HepG2 cells. (b) Venn diagram showing overlap of differentially expressed genes identified in total RNAs and chromatin associated RNAs by CAGE-seq analysis upon AGO1 knockdown. (c) Bar-plot showing number significantly (FDR 0.05) differentially expressed genes (up regulated and down-regulated) in each condition (total RNAs and chromatin bound RNAs). (d) Number of up-regulated and down-regulated miRNAs (P-value 0.01) identified by small RNA-seq in AGO1-KD. (e) Pie chart showing integration of AGO1-KD CAGE-seq and deregulated miRNAs (small RNA-seq) by using miRTarBase. 5% of differentially expressed genes (CAGE-seq) are predicted target of deregulated miRNAs and 95% of DE genes are non-target. (f-i) Bar-plot showing differential expression of four selected miRNAs and their predicted target genes (CAGE-seq) in AGO1 knockdown cells.

**Supplemental Figure 3 AGO1 ChIP-seq validation and analysis in HepG2 Cells.** (a) Independent AGO1 ChIP experiments followed by qPCR were performed in HepG2 cells to validate ChIP-seq data on selected genes that either contain (positive target) or not contain (negative target) AGO1 peaks at their promoter regions. Bar-plot shows percent input (% Input) enrichment of AGO1 on target genes (mean ± SD of three independent experiments). The IgG was used as mock control. (b) AGO1 knockdown decreases occupancy of AGO1 at the target gene promoters. HepG2 cells were transfected with control siRNA (siCtrl) or siRNA against AGO1 (siAGO1) for 72 hrs. ChIP-qPCR analysis was performed to determine AGO1 occupancy at the selected promoters. Bar-plot shows fold enrichment over mock (± SD from 2 independent experiments). (c) Heatmap showing enrichment of AGO1 ChIP-seq peaks (as blue shades) at enhancers (E) and promoters (TSS) regions determined by overlapping ChIP-seq peaks with chromatin segmentation states of HepG2 cells predicted by combination of ChromHMM and Segway. (d) Pie chart representing the distribution of AGO1 ChIP-seq peaks relative to genes (located over UTRs, exons, introns, 2-kbp upstream regions [promoter-TSSs], 1-

kbp downstream regions [TTSs]). The term “non-coding” refers to the exons of non-coding transcripts. (e) Bar plot showing enrichment of Ago1 ChIP-seq peaks at TSS (<1-kb) and distal regions (5-50-kb from TSS). (f) Density plot of ChIP-seq signals (median) for AGO1, and histone marks from ENCODE ChIP-seq data including H3K4me1, H3K4me3, H3K9ac, and H3K27ac centered at AGO1 peak located in distal, proximal and promoter regions. AGO1 is enriched with histone marks linked to active enhancers and promoters.

**Supplemental Figure 4 Affinity purification and mass spectrometry analysis of e-AGO1 and e-AGO2 complexes.** (a) Sketch of tandem affinity purification procedure. HEK-293 cell lines stably expressing FLAG-HA epitope tagged e-AGO1 and e-AGO2 were established. After fractionation into soluble nuclear and chromatin the e-AGO1 and e-AGO2 protein complexes were purified by double affinity purification, separated on SDS-PAGE and then subjected to silver staining, mass spectrometry analysis, and western blotting. (b) Paraspeckle proteins identified by mass spectrometry analysis in e-AGO1 complexes together with the number of identified peptides are indicated.

**Supplemental Figure 5 *NEAT1* ChIRP-western blotting identifies AGO1 as interacting partner.** (a) Sketch of *NEAT1* ChIRP-Western blot procedure. (b) ChIRP was performed using *NEAT1* probes (odd and even) and negative control LacZ probes. Isolated RNAs were analyzed by qRT-PCR using specific primers for *NEAT1* and GAPDH (negative target). Bar-chart shows successful *NEAT1* lncRNA retrieval (%) with both odd and even *NEAT1* probes (± SD from 3 experiments). (c) Proteins retrieved by *NEAT1* lncRNA and negative control probe along with input protein are visualized by silver staining. (d) Immunoblotting of *NEAT1* ChIRP-enriched proteins (isolated with odd and even pool) using specific antibodies to detect NONO, SFPQ, AGO1, AGO2, and

Tubulin. (e) Western blot for some essential paraspeckle proteins (NONO, SFPQ, HNRNPK and PSPC1) retrieved with *NEAT1* ChIRP in control and AGO1 knockdown cells. Results show a decrease in the level of NONO, SFPQ, and HNRNPK upon AGO1 depletion. (f) Western blot shows similar level of paraspeckle proteins (NONO, SFPQ, HNRNPK and PSPC1) in control and AGO1 knockdown cells. Tubulin was used as a loading control.

**Supplemental Figure 6 Sequencing statistics and quality control of Hi-C Library determined by HiC-Pro pipeline.** (a, b) The mapping and Hi-C library statistics for both replicates of siCtrl and siAGO1 cells. HiC-Pro quality control report for replicate2 of siCtrl and siAGO1 cells showing (c) statistics of read pairs alignment on restriction fragments and (d) fractions of duplicated reads and contact types. (e) Principal Component Analysis (PCA) plot showing the clustering of replicates for siCtrl and siAGO1 cells.

**Supplemental Figure 7 Differential interactions between control and AGO1-depleted cell.** (a) Differential interaction (siAGO1 versus siCtrl) heatmap for all chromosomes (1Mb), showing up-interacting regions in red and down- interacting bins in blue color. (b-c) Circos plots showing significant differentially interacting bins (FDR 0.01) at genome-wide level for (b) inter-chromosomal and (c) intra-chromosomal interactions. The outer layer represents human chromosomes in different colors.

**Figure 8 Changes in chromatin compartments and TADs in AGO1 depleted cells.** (a) First eigenvector values of chromosome 1 used to define A/B chromatin compartments, positive values represent open A-type (red) and negative values represent closed B-type (blue) compartments (top panel siCtrl and lower panel siAGO1 cells). Dotted lines delimit compartment borders. Expressed genes represent all genes identified by CAGE-seq both non-significant (genes with a basemean expression higher than 20 but not up-regulated upon AGO1 knockdown) and significant genes (with a basemean expression higher than 20 and being up-regulated for siAGO1 and down-regulated for siCtrl cells). (b) Box-plot showing average size (kb) of TADs identified in siCtrl and siAGO1 cells, at both 40-kb and 100-kb resolution (t−test p-value < 0.05). (c) Pie chart showing the percentage of AGO1 ChIP-seq peaks localized at TAD boundaries or not. (d) Venn diagram showing the number of overlapping TAD boundaries between siCtrl and siAGO1 cells.

**Supplemental Figure 9 Validation of expression changes for selected genes within differentially interacting loci.** Relative expression of selected example genes located within (a-b) gain or (c-d) decrease interacting loci was validated by (a and c) qRT-PCR. Values were normalized relative to the geometric mean of 18S rRNA and GAPDH mRNA expression levels. Error bars indicate mean ± s.e.m from 2 independent experiments. (b and d) Bar-plot shows the fold change values (from AGO1 KD CAGE-seq data for comparison) of genes located within (b) increase or (d) decrease interacting bins.

**Supplemental Figure 10 Quality control of Hi-C library in WT and *NEAT1*-KO HAP1 cell lines.** (a) Visualization of paraspeckles by *NEAT1* RNA-FISH and SFPQ immunofluorescence in wild type (WT) and *NEAT1* knockout HAP1 cells. (b-c) The mapping and Hi-C library statistics for both replicates of WT and *NEAT1*-KO HAP1 cells. (d-e) Genome-wide normalized Hi-C interaction heatmap at 1Mb resolution of (d) WT-HAP1 (e) *NEAT1*-KO HAP1 cell lines. Chromosomes are represented from top left to bottom right in order of chr1, chr2…chr22, chrX & ChrY. The color scale represents interaction frequencies.

#### Supplemental_Fig_S1

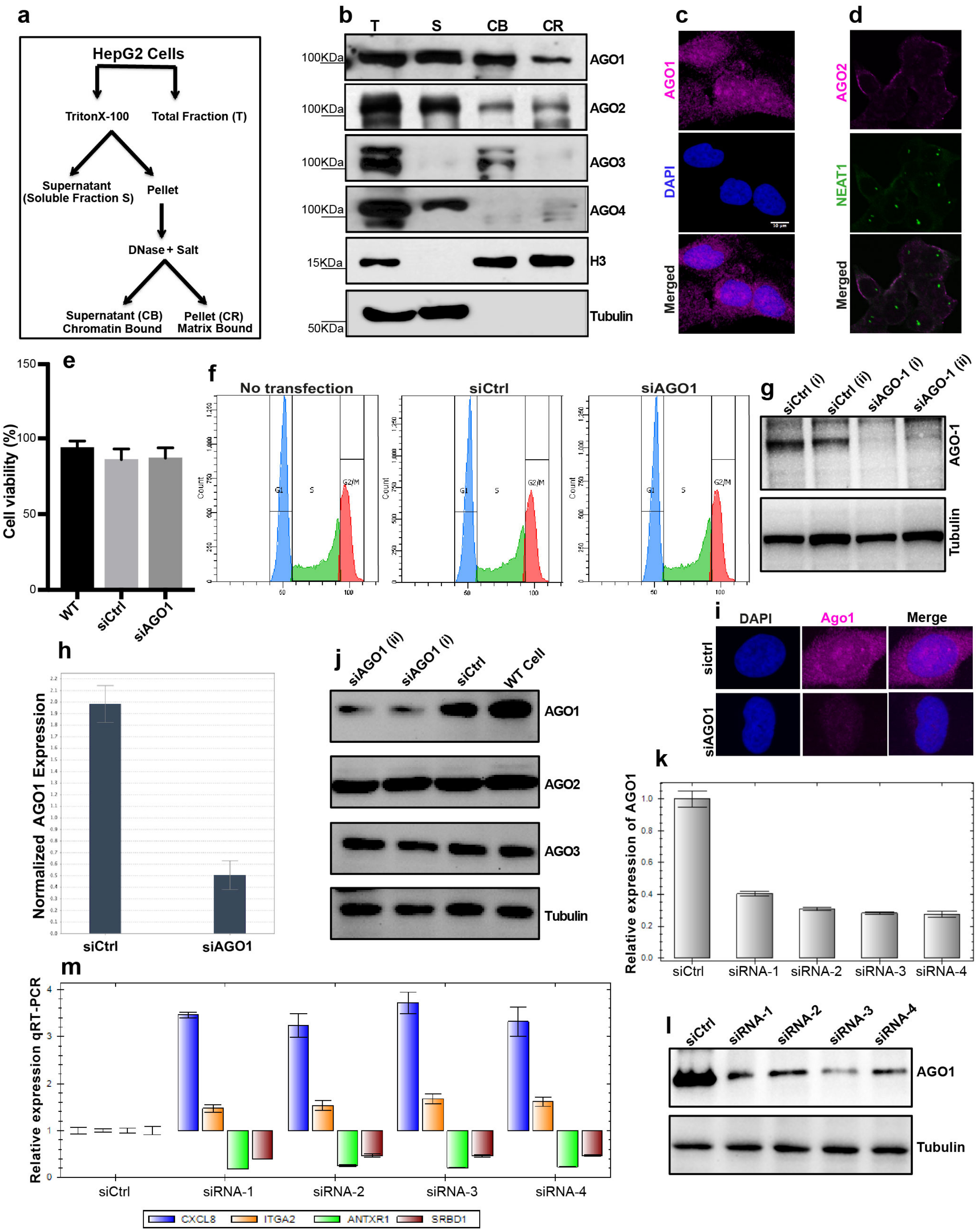

### Supplemental_Fig_S2

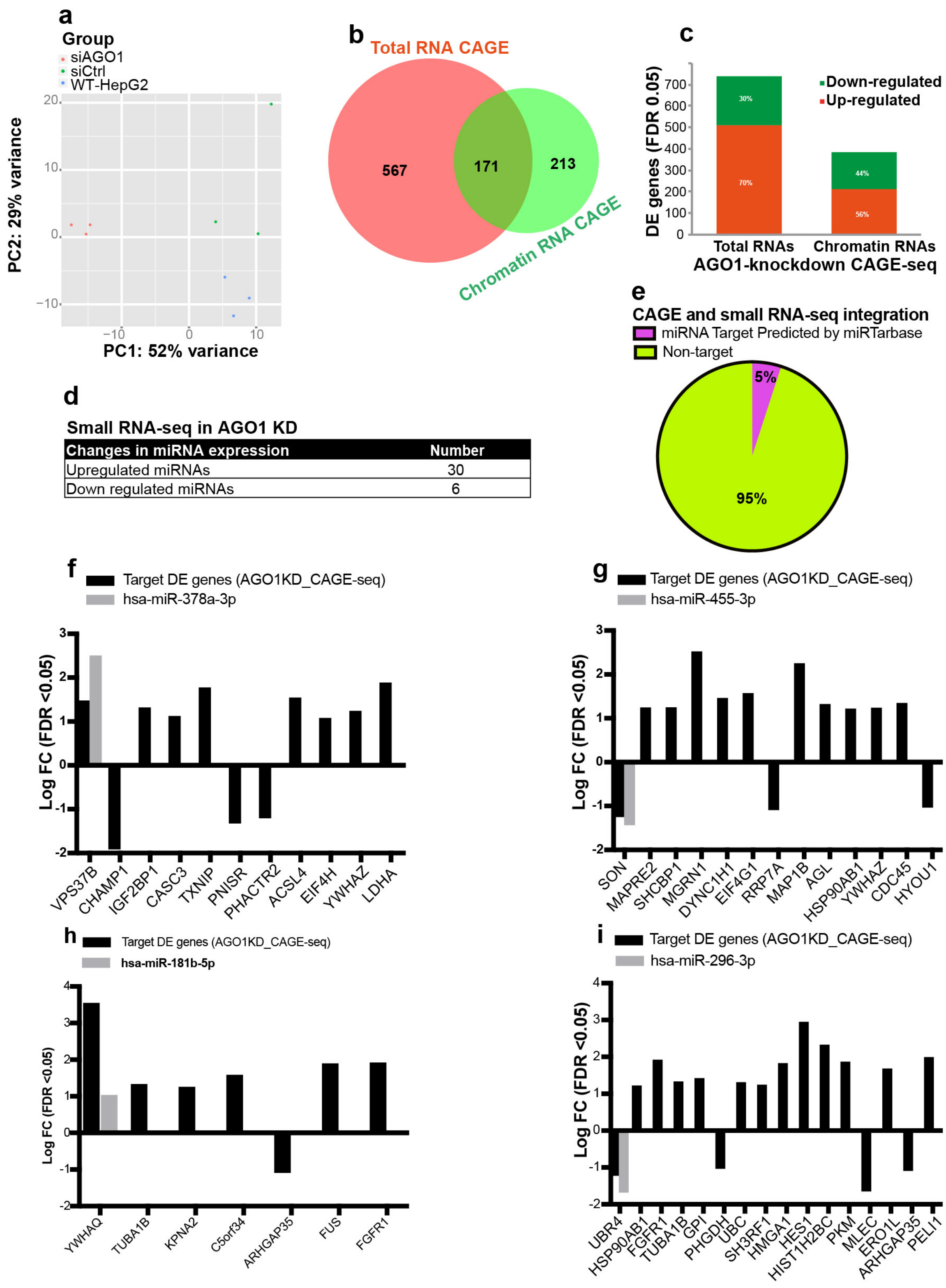

#### Supplemental_Fig_S3

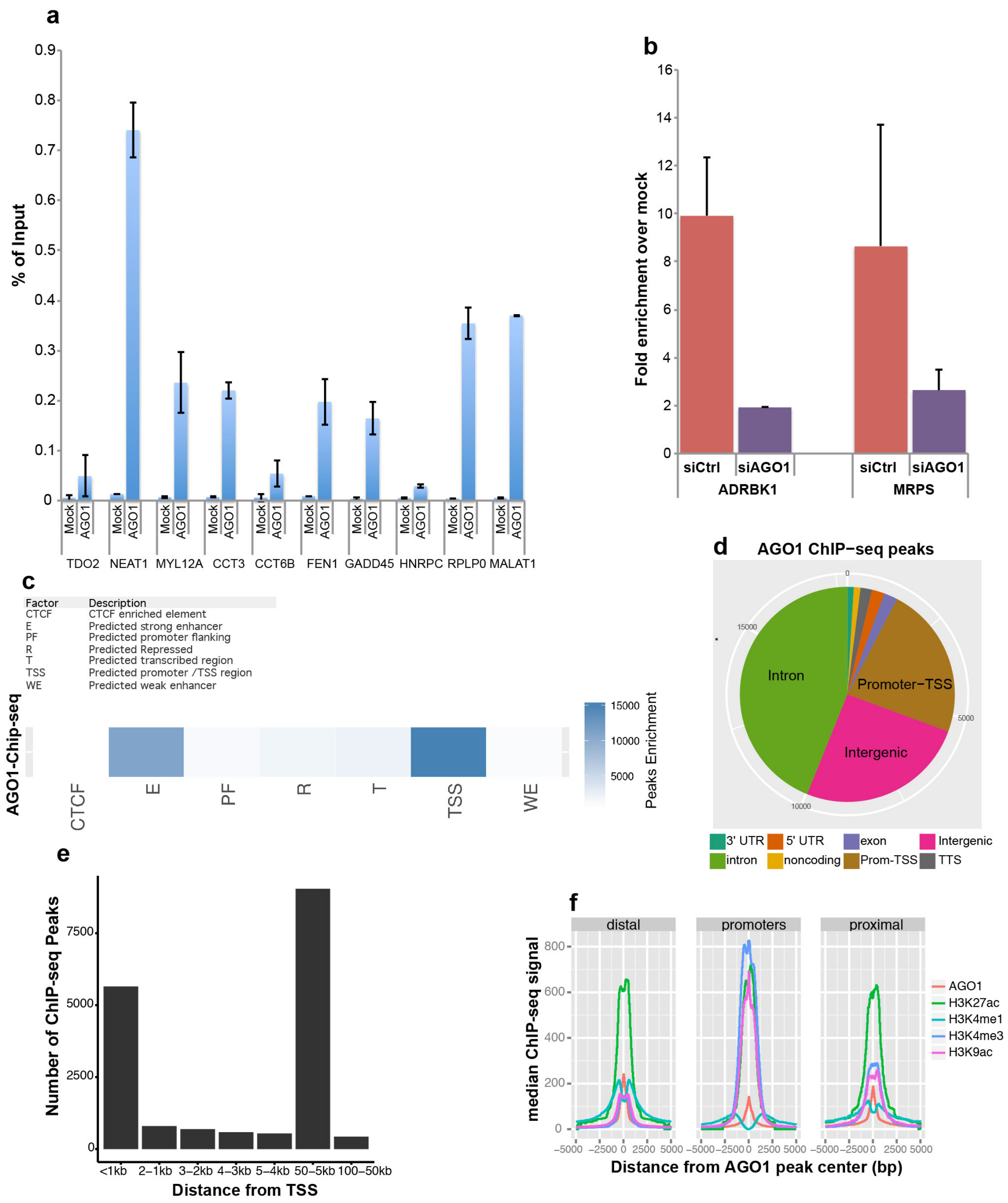

### Supplemental_Fig_S4

**a**

**Tandem Affinity Purification of AGO1/2 Complex**

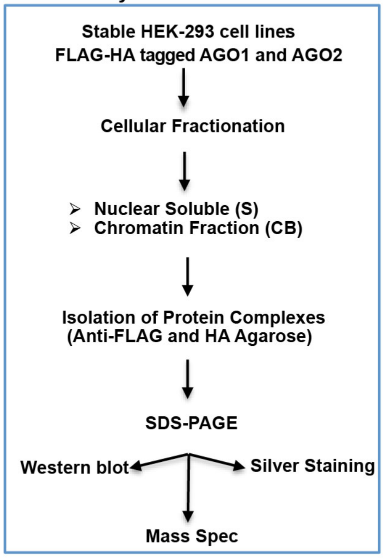

**b**

**Paraspeckle components associated with AGO1 complex**

| Protein | Accession # | MW (KDa) | Unique peptides identified | |
|---|---|---|---|---|
| | | | Nuclear soluble | Chromatin |
| FUS | P35637 | 53 | 5 | 8 |
| TARDBP | Q13148 | 45 | 6 | 10 |
| RBM14 | Q96PK6 | 69 | 5 | 7 |
| SFPQ/PSF | P23246 | 76 | 4 | 6 |
| HNRPF | P52597 | 45 | 5 | 6 |
| HNRPU | Q00839 | 90 | 4 | 7 |
| HNRPK | P61978 | 51 | 4 | 8 |
| HNRNPUL1 | Q9BUJ2 | 96 | 4 | 6 |
| HNRNPM | P52272 | 78 | 11 | 4 |
| HNRNPH3 | P31942 | 49 | 4 | 4 |

### Supplemental_Fig_S5

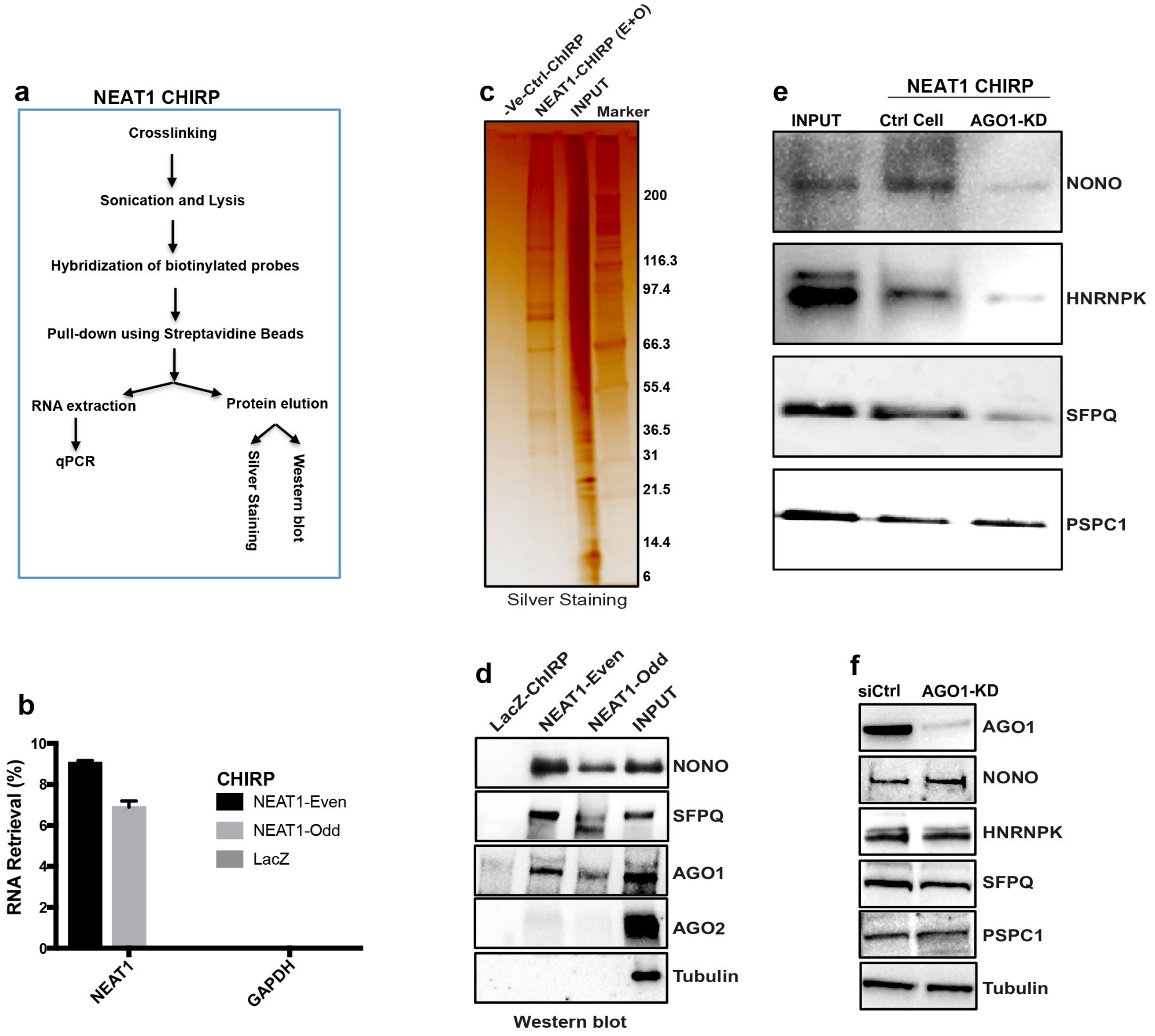

### Supplemental_Fig_S6

**a**

| Mapping Statistics | | | | | | | | | | |
|---|---|---|---|---|---|---|---|---|---|---|
| Sample Name | Individual Reads | | | | | | Paired Statistics | | | |
| | Total R1 reads | R1 Mapped | R1 Unmapped | Total R2 Reads | R2 mapped | R2 Unmapped | Total | Uniquely Mapped in pairs | Multimap in pairs | Pairs with singleton | Unmapped pairs |
| siCtrl- Rep1 | 106,133,981 | 88,924,057 | 17,209,924 | 106,133,981 | 79,924,030 | 26,209,951 | 106,133,981 | 65,677,552 | 8,065,750 | 21,361,483 | 11,029,196 |
| siCtrl-Rep2 | 131,504,936 | 109,496,402 | 22,008,534 | 131,504,936 | 100,618,521 | 30,886,415 | 131,504,936 | 82,640,403 | 10,114,423 | 24,605,271 | 14,144,839 |
| siAgo1-Rep1 | 121,575,943 | 109,188,154 | 12,387,789 | 121,575,943 | 101,649,904 | 19,926,039 | 121,575,943 | 87,504,634 | 9,157,825 | 17,513,140 | 7,400,344 |
| siAgo1-Rep2 | 113,258,385 | 101,302,918 | 11,955,467 | 113,258,385 | 89,262,278 | 23,996,107 | 113,258,385 | 76,036,240 | 8,458,936 | 21,574,844 | 7,188,365 |

**b**

| HiC Library Statistics | | | | | |
|---|---|---|---|---|---|
| Sample Name | Valid Interactions | Dangling_end_pairs | ReligationPairs | Self_cycle_pairs | ErrorPairs |
| siCtrl-Rep1 | 41,895,760 | 18,075,625 | 4,827,292 | 822,027 | 56,848 |
| siCtrl-Rep2 | 52,975,576 | 22,386,746 | 6,126,813 | 1,059,929 | 91,339 |
| siAgo1-Rep1 | 51,447,511 | 26,041,798 | 9,051,410 | 906,769 | 57,146 |
| siAgo1-Rep2 | 44,279,871 | 23,062,795 | 7,838,104 | 809,170 | 46,300 |

**c** Statistics of Read Pairs Alignment

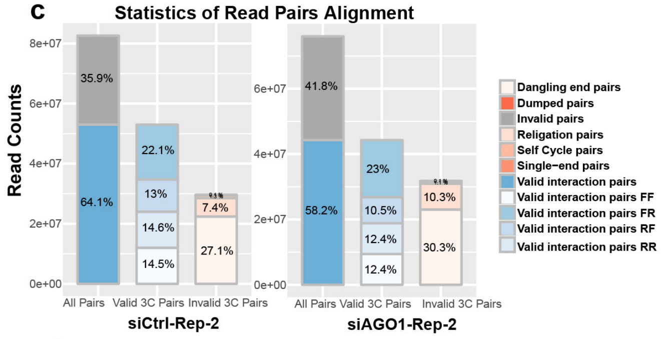

**d** Valid Pairs − duplicates and contact ranges

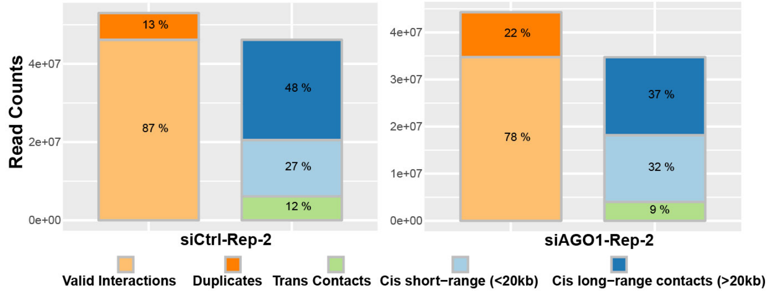

**e**

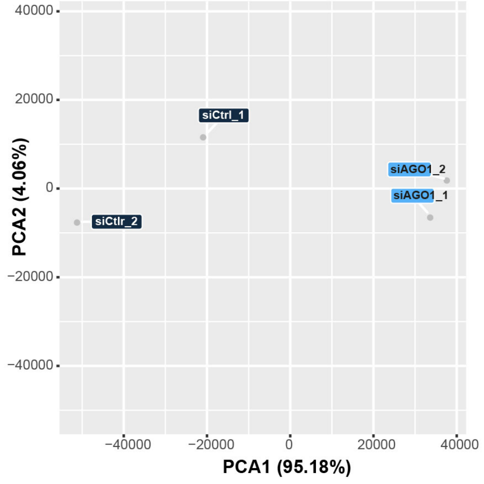

#### Supplemental_Fig_S7

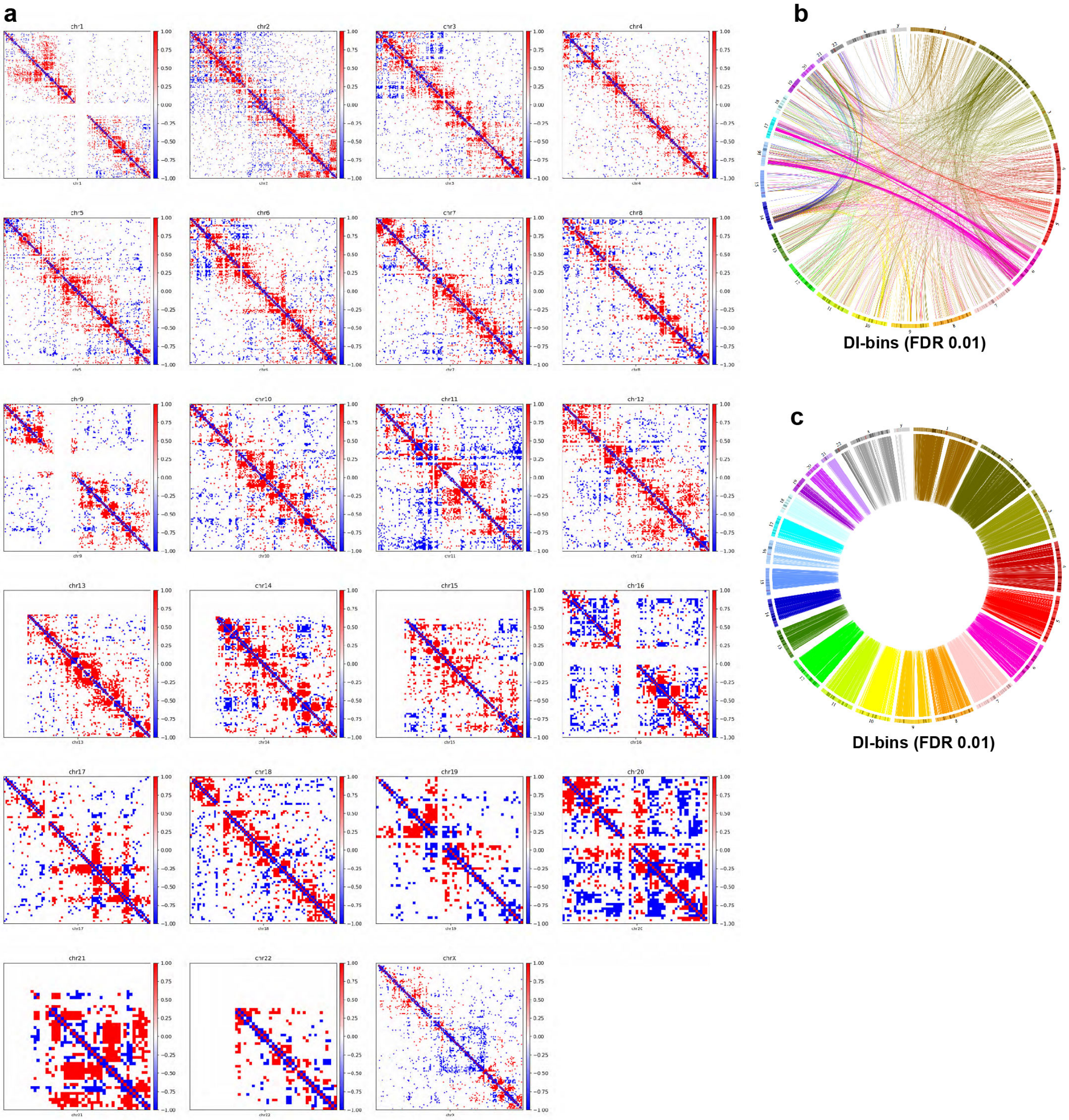

### Supplemental_Fig_S8

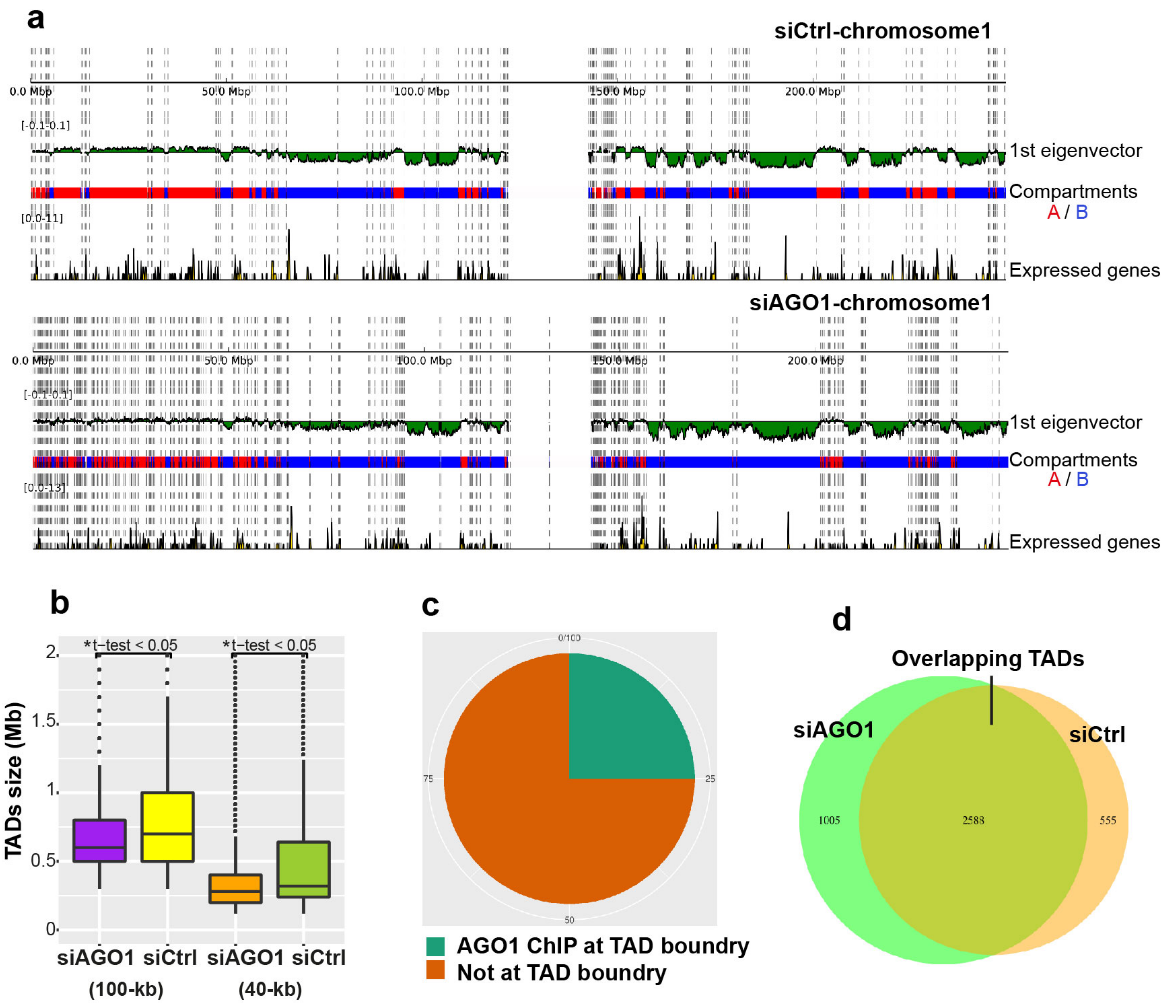

**Supplemental_Fig_S9**

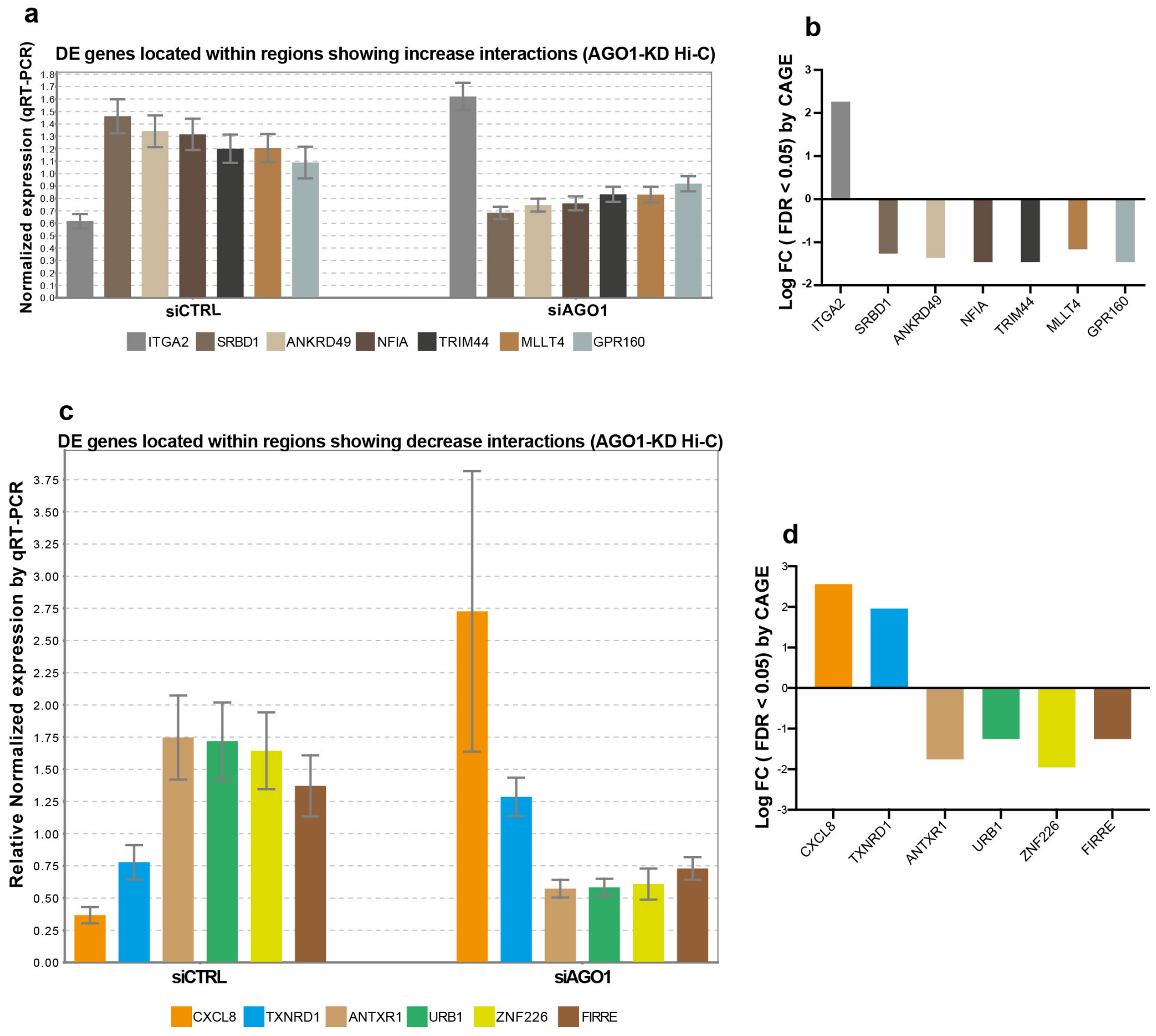

#### Supplemental_Fig_S10

**a**

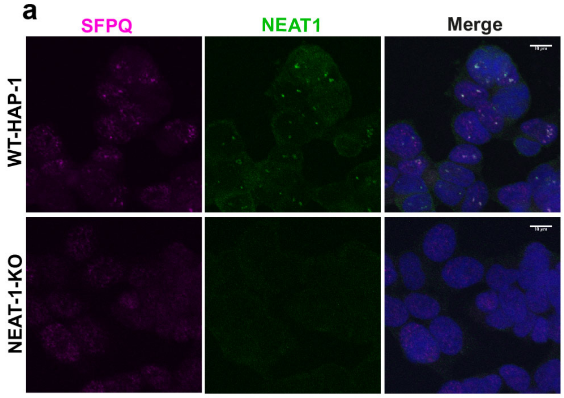

**b**

| Mapping statistic | | | | | | | |
|---|---|---|---|---|---|---|---|
| **Sample name** | **Total_pairs** | **Unmapped_pairs** | **Low_qual_pairs** | **Unique_paired** | **Multiple_pairs** | **Pairs_with_singleton** | **Reported_pairs** |
| **HAP1_WT-rep1** | 175645821 | 3420188 | 0 | 130892030 | 16149809 | 25183794 | 130892030 |
| **HAP1_WT-rep2** | 203108031 | 4439866 | 0 | 142744872 | 16986047 | 38937246 | 142744872 |
| **HAP1_NEAT1_KO-rep1** | 225644613 | 6167510 | 0 | 163035680 | 19502411 | 36939012 | 163035680 |
| **HAP1_NEAT1_KO-rep2** | 288771596 | 6524783 | 0 | 212890114 | 23895923 | 45460776 | 212890114 |

**c**

| Hi-C statistic | | | | | |
|---|---|---|---|---|---|
| **Sample name** | **Valid_interaction** | **Dangling_end_pairs** | **Religation_pairs** | **Self_Cycle_pairs** | **Dumped_pairs** |
| **HAP1_WT-rep1** | 96645607 | 11919658 | 20663212 | 1625320 | 38233 |
| **HAP1_WT-rep2** | 103255641 | 12184991 | 25231942 | 1988791 | 83507 |
| **HAP1_NEAT1_KO-rep1** | 96677206 | 32399882 | 32342936 | 1568807 | 46849 |
| **HAP1_NEAT1_KO-rep2** | 170869331 | 23493935 | 15686311 | 2677382 | 163155 |

**d**

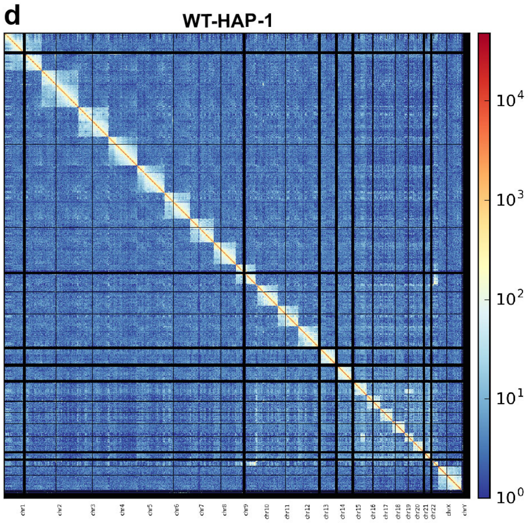

**e**

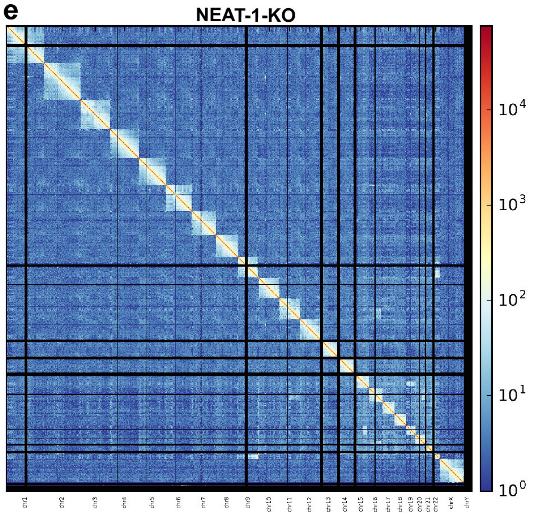
